# Dynamic evolution of euchromatic satellites on the X chromosome in *Drosophila melanogaster* and the *simulans* clade

**DOI:** 10.1101/846238

**Authors:** J.S. Sproul, D.E. Khost, D.G. Eickbush, S. Negm, X. Wei, I. Wong, A.M. Larracuente

## Abstract

Satellite DNAs (satDNAs) are among the most dynamically evolving components of eukaryotic genomes and play important roles in genome regulation, genome evolution, and speciation. Despite their abundance and functional impact, we know little about the evolutionary dynamics and molecular mechanisms that shape satDNA distributions in genomes. Here we use high-quality genome assemblies to study evolutionary dynamics of two complex satDNAs, *Rsp-like* and *1.688* gm/cm^3^, in *Drosophila melanogaster* and its three nearest relatives in the *simulans* clade. We show that large blocks of these repeats are highly dynamic in the heterochromatin, where their genomic location varies across species. We discovered that small blocks of satDNA that are abundant in X chromosome euchromatin are similarly dynamic, with repeats changing in abundance, location, and composition among species. We detail the proliferation of a rare satellite (*Rsp-like*) across the X chromosome in *D. simulans* and *D. mauritiana. Rsp-like* spreads by inserting into existing clusters of the older, more abundant *1.688* satellite, in events that were likely facilitated by microhomology-mediated repair pathways. We show that *Rsp-like* is abundant on extrachromosomal circular DNA in *D. simulans*, which may have contributed to its dynamic evolution. Intralocus satDNA expansions via unequal exchange and the movement of higher-order repeats also contribute to the fluidity of the repeat landscape. We find evidence that euchromatic satDNA repeats experience cycles of proliferation and diversification somewhat analogous to bursts of transposable element proliferation. Our study lays a foundation for mechanistic studies of satDNA proliferation and the functional and evolutionary consequences of satDNA movement.

## INTRODUCTION

Eukaryotic genomes are replete with large blocks of tandemly repeated DNA sequences. Named for their distinct “satellite” bands on cesium chloride density gradients (Kit 1961; Sueoka 1961; Szybalski 1968), these so-called satellite DNAs (satDNA) can comprise large fractions of eukaryotic genomes (Britten and Kohne 1968; Yunis and Yasmineh 1971). SatDNAs are a major component of heterochromatin—satDNAs accumulate in megabase-length blocks near centromeres and telomeres in many organisms (Charlesworth, et al. 1986; Charlesworth, et al. 1994). The location, abundance, and sequence of these heterochromatic satDNAs can turnover rapidly (Yunis and Yasmineh 1971; Ugarkovic and Plohl 2002) creating divergent repeat profiles between species (Strachan, et al. 1982). SatDNAs can be involved in intragenomic conflicts over transmission through the germline, as the driving centromeres that cheat female meiosis *(e.g.,* centromere drive, (Henikoff, et al. 2001), or targets of the sperm killers that cheat male meiosis (*e.g.,*Larracuente 2014; Courret, et al. 2019). These conflicts may fuel satDNA evolution. Changes in satDNA are expected to have broad evolutionary consequences due to their roles in diverse processes including chromatin packaging (Blattes, et al. 2006) and chromosome segregation (Dernburg, et al. 1996). For example, variation in satDNA can impact centromere location and stability (Aldrup-MacDonald, et al. 2016), meiotic drive systems (Fishman and Willis 2005; Fishman and Saunders 2008; Lindholm, et al. 2016), hybrid incompatibilities (Ferree and Barbash 2009), and genome evolution (Britten and Kohne 1968; Hartl 2000; Bosco, et al. 2007).

Small blocks of tandem repeats also occur in euchromatic regions of genomes (*e.g.*, Feliciello, et al. 2014; Ruiz-Ruano, et al. 2016) and are particularly enriched on Drosophila X chromosomes (Waring and Pollack 1987; DiBartolomeis, et al. 1992; Kuhn, et al. 2012; Gallach 2014). Some euchromatic X-linked repeats have sequence similarity to the large blocks of heterochromatic satDNAs *(e.g.,* Waring and Pollack 1987; DiBartolomeis, et al. 1992; Kuhn, et al. 2012) suggesting they could be a continual source of euchromatic repeats. Studies suggest these euchromatic repeats may play roles in gene regulation by acting as “evolutionary tuning knobs” (King, et al. 1997), regulating chromatin (Brajkovic, et al. 2012; Feliciello, et al. 2015), and facilitating X chromosome recognition/dosage compensation (Waring and Pollack 1987; Kuhn, et al. 2012; Lundberg, et al. 2013; Menon, et al. 2014; Lucchesi and Kuroda 2015; Joshi and Meller 2017; Deshpande and Meller 2018; Kim, et al. 2018).

Much of the species-level variation in satDNA arises through movement and divergence of an ancestral “library” of satellites inherited through common decent (Fry and Salser 1977). Unequal exchange between different repeats within a tandem array leads to expansions and contractions of repeats at a locus (Smith 1976), and along with gene conversion cause the homogenization of repeated sequences. This homogenization can occur both within repeat arrays *(e.g.,* Schlotterer and Tautz 1994) and between repeats on different chromosomes, causing repeat divergence between species (reviewed in Dover 1982). These processes result in the concerted evolution (Dover 1994) of satDNAs (Strachan, et al. 1982) and multicopy gene families like rDNA and histones (Coen, et al. 1982), leading to species-specific repeat profiles. Novel satDNAs can arise within a species from the amplification of unique sequences through replication slippage (Levinson and Gutman 1987; Schlotterer and Tautz 1992), unequal exchange, rolling circle replication (Britten and Kohne 1968; Southern 1970; Lohe and Brutlag 1987; Walsh 1987), and transposable element (TE) activity (Dias, et al. 2014; McGurk and Barbash 2018; Vondrak, et al. 2019). Recombination involving satDNA can cause local rearrangements or large-scale structural rearrangements such as chromosomal translocations (Richardson and Jasin 2000; Lieber, et al. 2006). Intra-chromatid recombination events give rise to extrachromosomal circular DNAs (eccDNAs) that are common across eukaryotic organisms (Cohen, et al. 1999; Cohen, et al. 2003; Cohen, et al. 2006; Zellinger and Riha 2007; Navratilova, et al. 2008; Cohen and Segal 2009; Paulsen, et al. 2018) and may contribute to the rapidly changing repeat landscape across genomes.

We have limited resolution on the evolutionary dynamics and molecular mechanisms that drive the rapid turnover of satDNA and its distribution in genomes. This lack of resolution is, in part, due to the challenges that repetitive DNA presents to sequence-based and molecular biology approaches. Here we characterize patterns and mechanisms underlying the evolution of complex satellites over short evolutionary time scales in *Drosophila melanogaster* and the closely related species in the *simulans* clade, *D. mauritiana, D. sechellia,* and *D. simulans.* We focus on two abundant satellite repeat families: *1.688 gm/cm3* and *Rsp-like. 1.688 g/cm3* (hereafter called *1.688)* is a family of several related repeats named after their monomer lengths, including *260bp, 353bp, 356bp, 359bp,* and *360bp* (Losada and Villasante 1996; Abad, et al. 2000). *Rsp-like* is a 160-bp repeat named for its similarity to the 120-bp *Responder (Rsp)* satellite (Larracuente 2014). We studied broad-scale patterns using cytological and genomic approaches. By leveraging new reference genomes based on long single-molecule sequence reads (Chakraborty, et al., unpublished), we study the dynamics of these repeats at base-pair resolution across the X chromosome. We discovered the rapid spread of *Rsp-like* repeats to new locations across the X chromosome in *D. simulans* and *D. mauritiana.* We explored the mechanism of satDNA movement, including the potential role of interlocus gene conversion and eccDNA in facilitating the spread of satellites across long physical distances on the X chromosome. Revealing the processes that shape satDNA evolution over short time scales is a critical step toward understanding the functional and evolutionary consequences of repeat turnover.

## RESULTS

### Heterochromatic and euchromatic satDNA composition varies across species

Our analysis of mitotic chromosomes with fluorescence *in situ* hybridization shows that large heterochromatic blocks of *1.688* repeats are primarily X-linked in *D. melanogaster* and *D. sechellia* but are autosomal in *D. simulans* and *D. mauritiana* (Fig. S1). *D. melanogaster* also has two smaller blocks of *1.688* family repeats in the heterochromatin of chromosome 3 (Abad, et al. 2000). The distribution of the *Rsp-like* family is similarly dynamic in the heterochromatin: large blocks are X-linked in *D. simulans,* autosomal in *D. sechellia* (chromosome 2 and 3), and lacking in the heterochromatin of *D. mauritiana* and *D. melanogaster* (Larracuente 2014); Fig. S1). The *1.688* repeat family also exists in the euchromatin (Waring and Pollack 1987; DiBartolomeis, et al. 1992; Kuhn, et al. 2012; Gallach 2014), where they are over-represented on the X chromosome relative to the autosomes in these *Drosophila* species (Chakraborty, et al., unpublished).

We mapped euchromatic satDNA repeats at a fine scale across the X chromosome. We find that similar to *1.688, Rsp-like* repeats are also present in the X euchromatin (Figs. 1, S2–3). We describe the location of these repeats relative to their cytological divisions *(i.e.* cytobands) on *D. melanogaster* polytene chromosomes and hereafter use the terms ‘cytobands’, ‘clusters’, and ‘monomers’ as illustrated in Fig. 1a. Both satellites accumulate near the telomere (cytoband 1) and in the middle of the X chromosome but are uncommon from cytoband 15 to the centromere (Figs. 1, S3). We confirmed the euchromatic enrichment of these repeats using FISH on polytene chromosomes, where we see a high density of bands on the polytenized arm of the X chromosome in the *simulans* clade species *(e.g.,* representative FISH image; Fig. S2).

**Figure 1:**
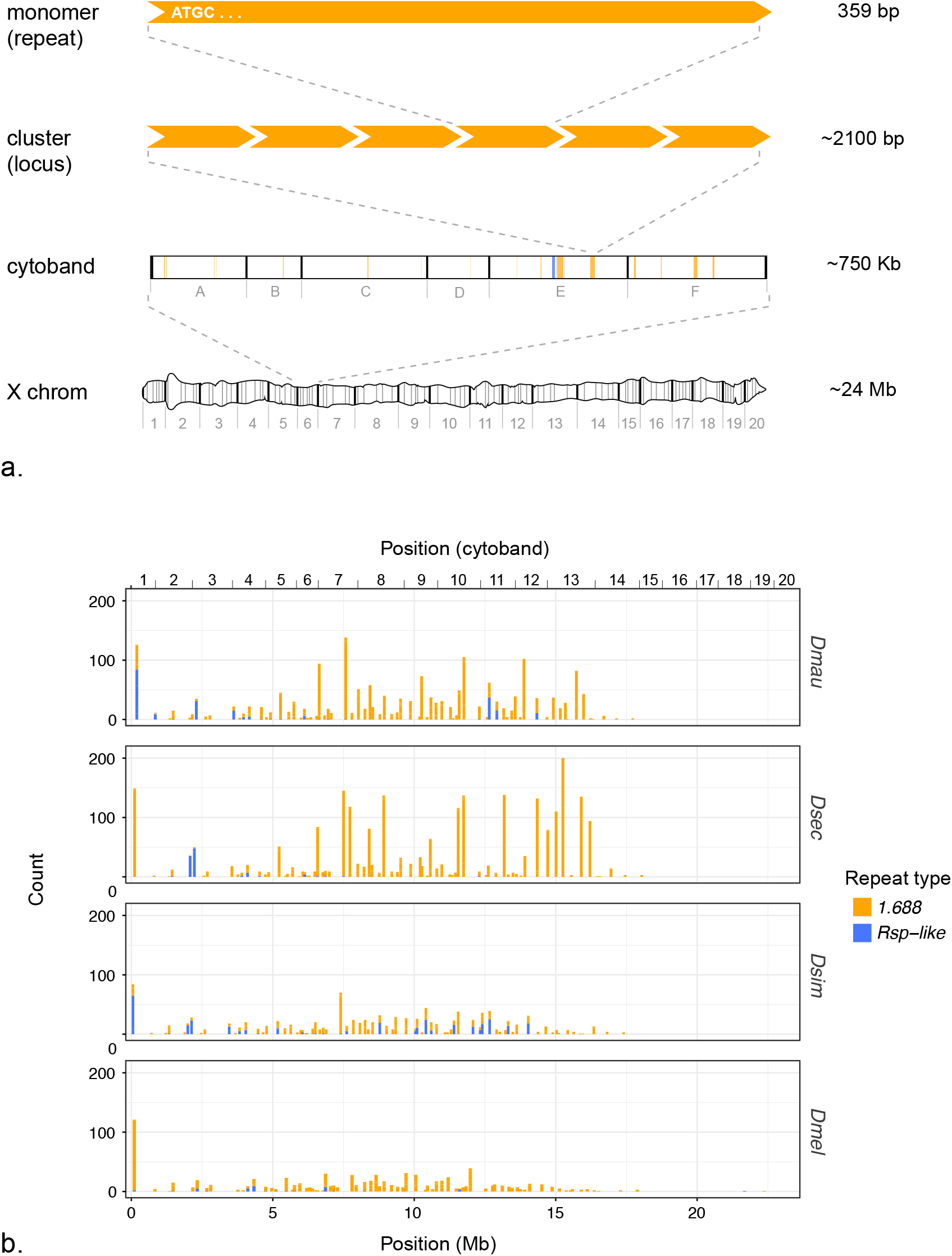
Euchromatic X-linked satellites are unevenly distributed across the X chromosome. (a.) A schematic illustrating terms frequently used in the text. We use ‘cytoband’ to reference large regions of the X chromosome that are defined by banding patterns in polytene chromosomes. We use ‘cluster’ to mean any distinct genomic locus containing the repeat of interest; typically >1 repeat. ‘Monomer’ refers to a single repeat unit; the example shown represents a *1.688* monomer. (b.) The x-axis shows position of *1.688* and *Rsp-like* satDNA clusters along the X chromosome. Each bar on the chart represents a cytological subdivision *(e.g.,* 1A, 1B, etc.) in which counts of all repeats are pooled. The y-axis indicates the number of repeat copies *(i.e.,* monomers) within a subdivision.

The abundance of euchromatic complex satellite repeats shows a 3-fold variation among species. *D. sechellia* has the most euchromatic X-linked repeats (2588 annotations), followed by *D. mauritiana* (1390), *D. simulans* (1112) and *D. melanogaster* (849) (Table 1). The *D. sechellia* X chromosome assembly contains 19 gaps, six of which occur within satellite loci (Chakraborty, et al., unpublished), therefore the X-linked copy number represents a minimum estimate for this species.

**Table 1.** Summary of euchromatic satDNA cluster sizes on X chromosome. Total #: number of total repeats. # clust: total number of clusters at distinct loci. % N=1: percentage of singletons (clusters of a single repeat). % N<4: percentage of small clusters (less than four repeats).

| Species | Total#<br>*1.688* | # *1.688*<br>clust | %N=1<br>*1.688* | % N<4<br>*1.688* | # *Rsp-*<br>*like* | # *Rsp-*<br>*like* clust | % N=1<br>*Rsp-like* | % N<4<br>*Rsp-like* |
| --- | --- | --- | --- | --- | --- | --- | --- | --- |
| *D. mauritiana* | 1165 | 325 | 24.00 | 68.31 | 225 | 26 | 30.77 | 34.62 |
| *D. sechellia* | 2486 | 308 | 33.44 | 82.14 | 102 | 12 | 50.00 | 58.33 |
| *D. simulans* | 786 | 324 | 31.17 | 89.20 | 326 | 38 | 18.42 | 34.21 |
| *D. melanogaster* | 808 | 274 | 33.94 | 83.94 | 41 | 19 | 73.68 | 78.95 |

Across all species, *1.688* is more abundant than *Rsp-like,* both in terms of total repeats *(i.e.,* the number of euchromatic repeat monomers annotated in our assemblies, Fig. 1a) and the number of clusters *(i.e.,* the number of distinct genomic loci containing repeats, Fig. 1a). Single-monomer clusters exist in both satDNA types; they represent ~30% of all *1.688* clusters and ~43% (~33% if *D. melanogaster* is excluded) of all *Rsp-like* clusters (Table 1, Fig. S4). These single-monomer clusters are considered “dead” as they cannot undergo unequal exchange and expand (Dover 1982; Langley, et al. 1988; Charlesworth, et al. 1994). The majority of the remaining *1.688* clusters are also small (*i.e*., contain 2-3 repeats) while the majority of the remaining *Rsp-like* clusters are larger (i.e., contain ≥4 repeats; Table 1, Fig. S4).

Both the number of total repeats and the number of clusters for each satellite varies among species. *Rsp-like* shows an 8-fold difference in total repeat number and a 3-fold difference in number of clusters across species, with *D. simulans* and *D. mauritiana* having more total repeats as well as more clusters than *D. sechellia* and *D. melanogaster* (Table 1, Fig. S4). In *D. simulans* and *D. mauritiana, Rsp-like* clusters have apparently spread to cytobands that lack such clusters in one, or both of the other species (*e.g.*, clusters at cytobands 7-12 in *D. simulans* and cytobands 11-12 in *D. mauritiana;* Figs. 1, S3). A relatively recent spread is consistent with *D. simulans* and *D. mauritiana* having a lower proportion of single repeat, or ‘dead’ clusters (18.4% and 30.8%, respectively) than the other species (Table 1). In *1.688, D. sechellia* shows as much as a 3-fold increase in total repeats despite having fewer *1.688* loci than the other *simulans* clade species, a pattern driven by a high number of large clusters in *D. sechellia* (16 clusters with ≥50 monomers), which are less common in other species (six clusters in *D. mauritiana*, one in both *D. simulans* and *D. melanogaster*; Table 1).

These patterns suggest dynamic turnover of satDNA repeat composition across the X chromosome euchromatin over short evolutionary time scales. While it is tempting to make a sweeping statement about a shift in repeat composition based on these numbers, it is difficult to systematically identify orthologous loci across the X chromosome to accurately quantify the turnover on a locus-by-locus basis. However, we can explore the dynamics of specific clusters for which synteny of unique flanking sequences strongly suggests orthology across species. One such representative cluster is embedded between two genes—*echinus* and *roX1—*at cytoband 3F (Fig. 2). In *D. melanogaster,* this cluster has only two *1.688* repeats, the first of which is truncated, plus an unannotated adjacent region that contains degenerated *1.688* sequence. *D. sechellia* also has *1.688* at this location, but the cluster is expanded relative to *D. melanogaster.* In contrast, both *Rsp-like* and *1.688* repeats are present at this locus in *D. mauritiana* and *D. simulans;* however, each species shows differences in repeat number of the respective satellites (Fig. 2).

**Figure 2:**
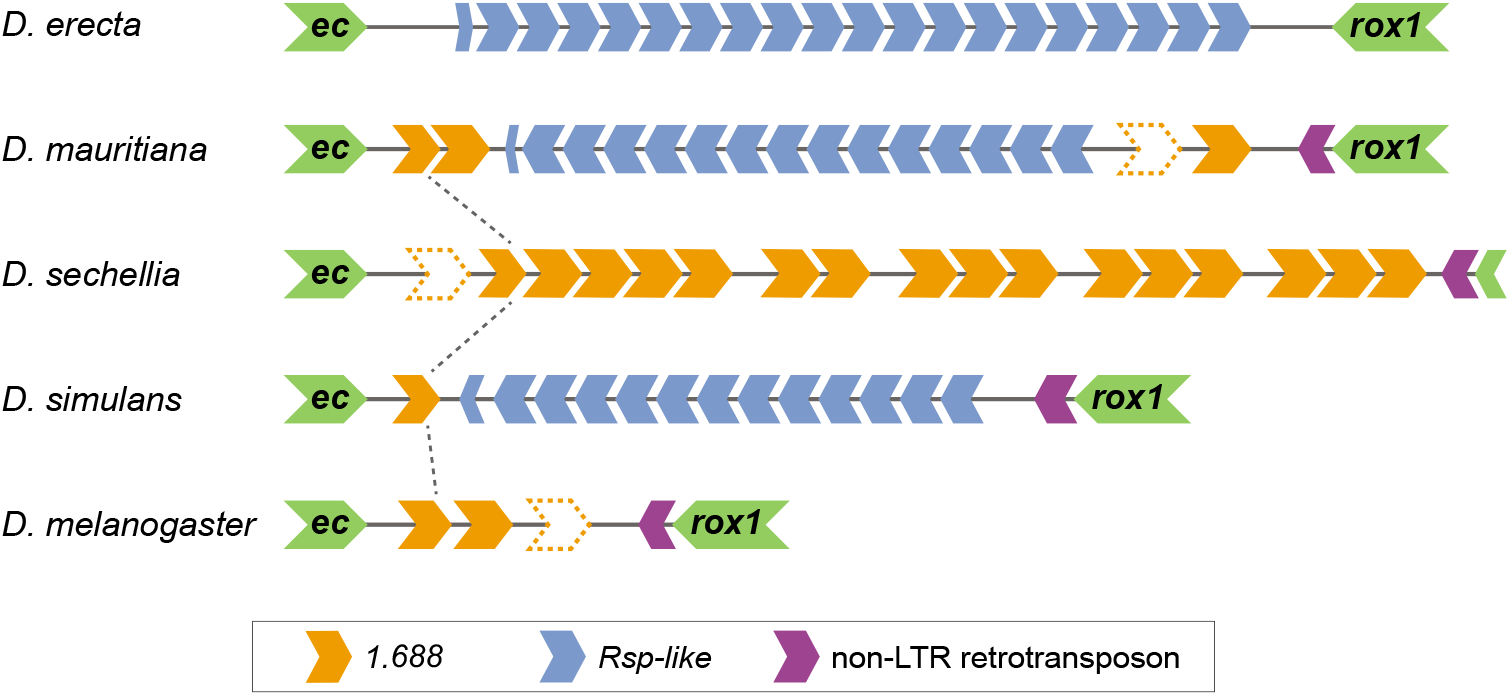
Organization of cytoband 3F repeat cluster. Schematic of 3F cluster in *D. melanogaster* and the simulans clade, as well as the outgroup species *D. erecta.* Cluster is flanked by two genes, *echinus* and *roX1* (green chevrons), with a TE insertion at the distal side of the locus (purple chevrons). Complex satellite monomers are indicated by blue *(Rsp-like)* or orange *(1.688)* chevrons. Chevrons with dotted outline indicate sequences that were not annotated, but were determined manually by BLAST to be highly degenerated satellite monomers. Black dotted lines between species indicate shared repeats.

The *Rsp-like* repeats in *D. mauritiana* and *D. simulans* are homogenized within each locus but are highly divergent between species. The major differences in X-linked satellite composition among species suggest that euchromatic satellites, like heterochromatic satellites, evolve dynamically over short evolutionary time scales.

### Recent proliferation of satDNA across the X euchromatin

Analysis of the nearest upstream and downstream genomic features relative to *1.688* and *Rsp-like* satellites showed that *Rsp-like* clusters have a non-random distribution, particularly in *D. simulans* and *D. mauritiana. Rsp-like* clusters are directly adjacent to, or interspersed with, *1.688* clusters in 82% of euchromatic X-linked clusters in *D. simulans* and in 62% of clusters in *D. mauritiana* (Table 2, Figs. S5–S6). Conversely, the *1.688* clusters do not seem to preferentially associate with *Rsp-like*, though they are often located near genes consistent with previous findings (Kuhn, et al. 2012); Figs. S5–S6).

**Table 2:** *Rsp-like* clusters associate with *1.688*. # *Rsp-like:* number of *Rsp-like* clusters on X chromosome. *#Rsp-like* / *1.688:* number of *Rsp-like* clusters (including singletons) that have *1.688* repeats within 100bp either upstream or downstream.

| Species | *Rsp-like* | *Rsp-like* / *I.688* | % *Rsp-like* / *I.688* |
| --- | --- | --- | --- |
| *D. mauritiana* | 26 | 16 | 62 |
| *D. sechellia* | 12 | 3 | 25 |
| *D. simulans* | 38 | 31 | 82 |
| *D. melanogaster* | 19 | 7 | 37 |

Examination of within-species and all-species phylogenetic trees of satellite repeats led to four major findings. (1) Heterochromatic repeats form clades that are generally separate from euchromatic repeats for both satellites in all species except *D. sechellia*, for which euchromatic and heterochromatic repeats are interspersed in both *1.688* and *Rsp-like* (Figs. S7–14). (2) *D. sechellia* and *D. mauritiana* (especially the former) show repeated evidence of intralocus expansion of repeats (Figs. S15–16). (3) *1.688* euchromatic repeats have a relatively old diversification history that largely pre-dates the speciation events that gave rise to the study species (Figs. 3–4, S7, S9, S11, S13, S15–16). This contrasts with *Rsp-like,* which shows evidence of relatively recent diversification, particularly in the *simulans* clade species (Figs. 3–4, S8, S10, S12, S14, S17–18). (4) *Rsp-like* repeats show evidence of two major expansions (Figs. 3–4, S8, S10, S12, S17–18), which encompass large physical distances across the X chromosome (*i.e*., ‘interlocus’ expansions) and mainly occurred independently in *D. simulans* and *D. mauritiana*. The latter two findings are discussed in additional detail in Supplemental Material.

**Figure 3.**
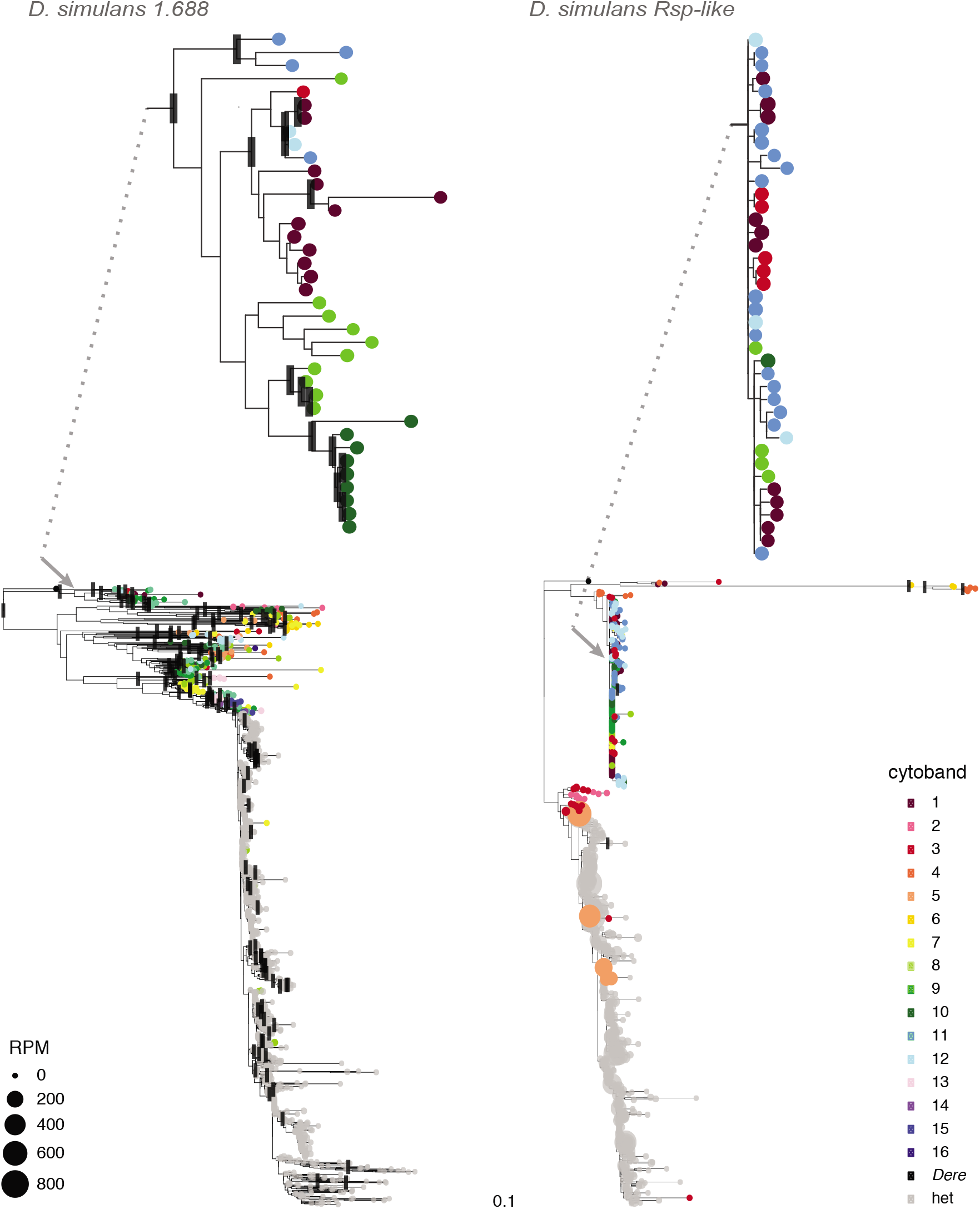
Comparison of phylogenetic patterns *1.688* and *Rsp-like* for *D. simulans.* Each terminal represents an individual repeat monomer from the X chromosome. Colored tip terminals indicate euchromatic repeats; gray tip terminals represent repeats from heterochromatic loci (defined as unassigned scaffolds in the assembly). Black rectangles indicate nodes with bootstrap support ≥ 90. Two regions in each tree are shown in greater detail to highlight differential phylogenetic patterns observed in euchromatic repeats of *1.688* and *Rsp-like*; arrows and dotted lines indicate relative position of enlarged regions in the tree. Branch lengths shown are proportional to divergence with both trees shown on the same relative scale. Sizes of the tips are scaled to reflect proportion of eccDNA reads mapping to a given variant, expressed as reads-per-million (RPM) (see eccDNA analysis). Maximum likelihood trees were inferred in RAxML with nodal support calculated following 100 bootstrap replicates.

**Figure 4:**
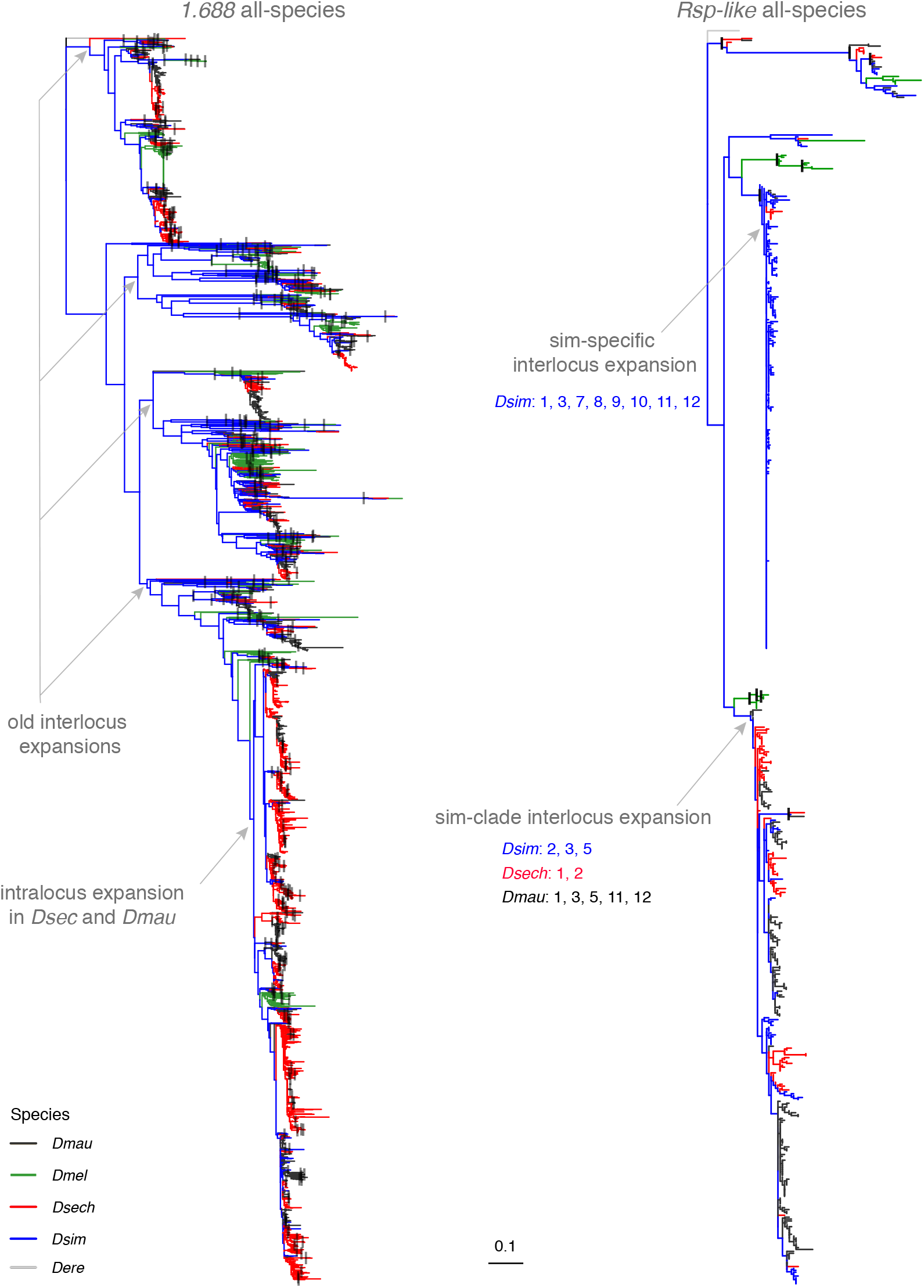
All-species maximum likelihood trees of euchromatic *1.688* and *Rsp-like.* Each terminal represents an individual repeat monomer. All monomers from clusters with ≥three repeats were included in the analysis. Species identity is indicated by branch color. Major inter and intralocus expansions of satellites discussed in the text are labeled with gray arrows. For interlocus expansions in *Rsp-like*, the species involved are listed along with cytological bands that are represented by monomers within the expansion. The outgroup (*D. erecta)* is indicated by gray branches. Black rectangles indicate nodes with bootstrap support ≥ 90. Maximum likelihood tree was inferred in RAxML with nodal support calculated following 100 bootstrap replicates. Branch length is shown proportional to relative divergence with both trees on the same relative scale. See Figures S15–18 for added detail as to genomic location of terminals.

### Mechanisms driving satellite DNA turnover in the euchromatin

How did the new *Rsp-like* clusters observed in *D. simulans* and *D. mauritiana (i.e.,* finding four of the previous section) arise? We found frequent co-localization of *Rsp-like* and *1.688* repeats in these species, which was surprising because these two repeats are unrelated at the sequence level. We therefore hypothesized that regions of microhomology could facilitate insertion of new *Rsp-like* repeats into pre-existing *1.688* clusters.

Our analysis of the *1.688/Rsp-like* junctions on each side of newly inserted *Rsp-like* clusters in *D. simulans* and *D. mauritiana* revealed multiple independent insertion events with shared signatures (Fig. 5). One prominent signature is that junctions between the *Rsp-like* and *1.688* sequences commonly occur at positions of microhomology. The same junction sequence is often shared between clusters at different locations across the X chromosome. We use the sequence of these microhomologies to define clusters of the same “type”: type 1 was found in *D. simulans* and types 2 and 3 were found in *D. mauritiana.* Because there are different *1.688* variants adjacent to both type 1 and 2 junctions (*e.g.*, compare Dsim10A and Dsim11E1, Fig. 5), we infer that five or more independent events have created the three junction types.

**Figure 5.**
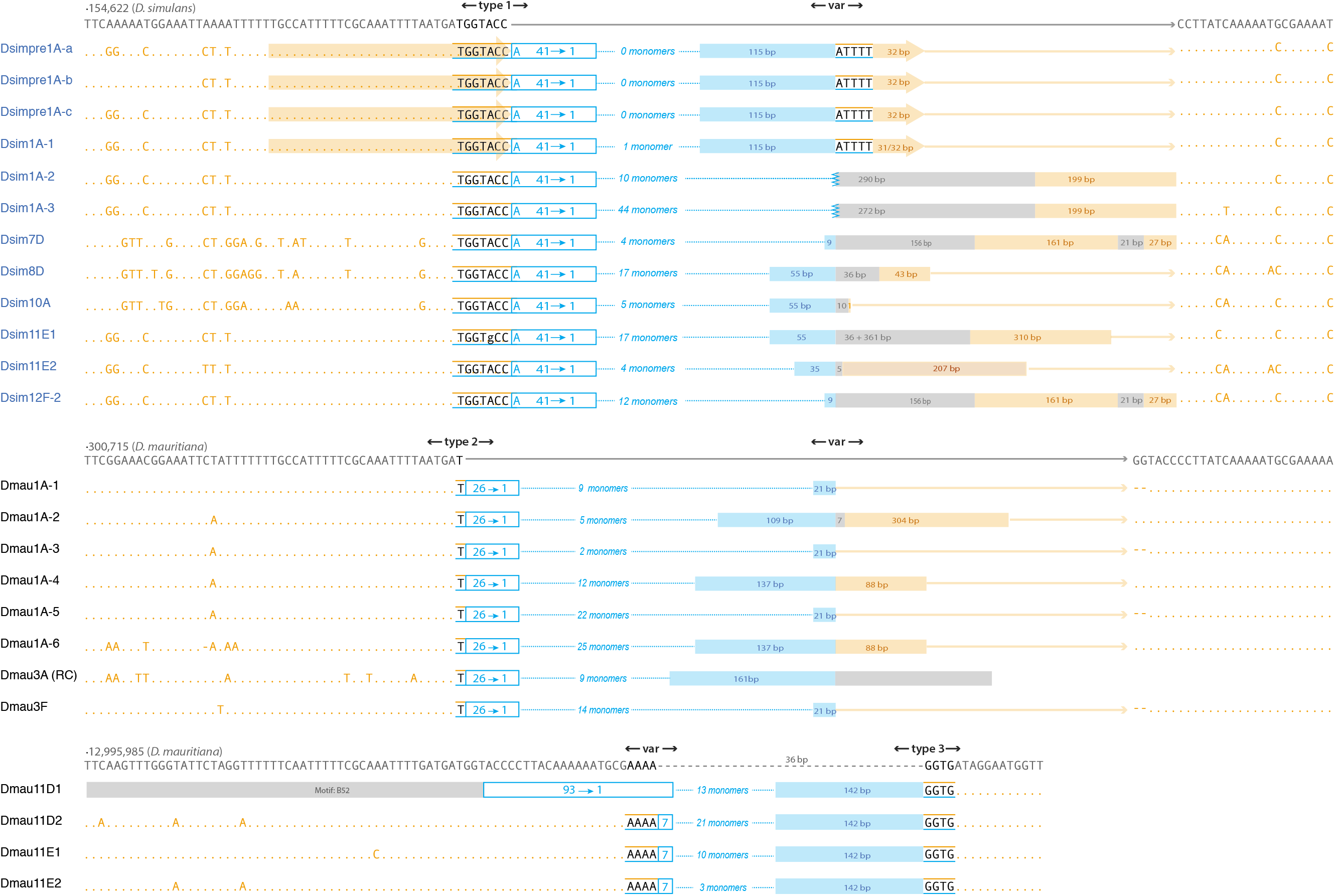
Junctions at new Rsp-like insertions in *D. simulans* and *D. mauritiana.* Junctions from a subset of the newer *Rsp-like* clusters (blue text/lines/boxes) are aligned and grouped into three types based on common signatures with nearby *1.688* monomers (orange text/lines/boxes). Type 1 is found in *D. simulans* while types 2 and 3 junctions are found in *D. mauritiana* (cytoband location of each cluster is indicated in the names at far left). Within each type, identical truncated *Rsp-like* monomers abut *1.688* at the same position in the *1.688* repeat monomer. In all three junction types, there is overlap between the two satellite sequences (black text) which, for at least the longer overlaps, potentially represents microhomology involved in the original insertion event. The second junction associated within and among these types is more variable (“var” in figure) with *Rsp-like* sequences abutting different positions of the *1.688* repeat or different unannotated sequences (gray boxes). The number of full length *Rsp-like* monomers as well as the lengths of truncated *Rsp-like* monomers, unannotated regions, and *1.688* sequences in this variable region are indicated for each cluster. Note that some clusters are nearly identical across this variable region *(e.g.,* Dsim7D and Dsim12F). The *1.688* sequences in the region that would be sequential to those sequences at the conserved junctions (dark gray text above each junction type is the sequence within a specific *1.688* monomer) are indicated at the far right. Orange arrows in the first four *D. simulans* clusters indicate a duplication of the *1.688* sequences at the two junctions.

In *D. simulans,* type 1 is the predominant junction and is observed in 19/31 *Rsp-like* clusters located near *1.688* repeats, 12 of which are diagrammed in Figure 5. The type 1 junction is associated with a 42/49 bp truncated *Rsp-like* monomer abutting *1.688* sequences. The transition between the two satellite types includes a 7-bp region of microhomology (‘TGGTACC’). Among these 12 *Rsp-like* clusters there are, however, at least 6 different junction sequences at the other end of the cluster. One of these variable junctions includes four clusters in which the sequences adjacent to *Rsp-like* are a duplication of the 32 bp (including the microhomology) of *1.688* sequences found at the type 1 junction. The remaining clusters have varying lengths of unannotated (5 bp to 290 bp) and *1.688* sequences (1 bp to 310 bp) in the variable region. All 19 *Rsp-like* insertions, which includes the clusters at 3F, 9D, 9F, 11C, 11D, 12C, and 12F-1 not diagrammed in Figure 5, are associated with a minor subset of *1.688* repeat variants comprising ~15% of the 787 monomers examined.

In *D. mauritiana*, type 2 clusters show a similar signature to *D. simulans* type 1 clusters: one side of the cluster shows a characteristic junction which is associated with a *Rsp-like* truncated monomer abutting *1.688* sequences, with the other side of the cluster showing more variable patterns. Interestingly, type 2 junctions occur at nearly the same position within the *1.688* monomer and in a similar subset of variants as the *D. simulans* type 1 junction, however, the position in *Rsp-like* monomers associated with the junction differs between the two species *(i.e.,* note 26/27 bp truncated monomers in *D. mauritiana* and 42/49 bp truncations in *D. simulans;* Fig. 5). The variable side of the cluster shows 4 different sequences associated with the junction. The most common variable junction occurs in four of the eight clusters and has a 2 bp deletion before continuing with the interrupted *1.688* repeat sequence. Likewise, the 4 new clusters in cytoband 11 of *D. mauritiana* show these junction signatures although unlike the type 1 and type 2 junctions, these type 3 junctions have a larger deletion (36 bp) in the associated *1.688* sequences.

The nature of the variable junctions (unannotated sequences/sequence variation in *1.688* repeat monomers) makes it difficult to determine whether insertion was facilitated by microhomology at these junctions. However, in two cases short runs of mononucleotides are present at the overlap between *1.688* and *Rsp-like* sequences. While non-homologous end joining (NHEJ) does not require, but can use, short stretches of microhomology (< 5 bp; Chang, et al. 2017), the multiple occurrences of microhomology including the 7 and 4 bp of microhomology observed in the type 1 and type 3 junctions, respectively, suggest that pathways employing microhomology-mediated end joining (MMEJ) facilitate *Rsp-like* insertions (Fig. 6a).

**Figure 6.**
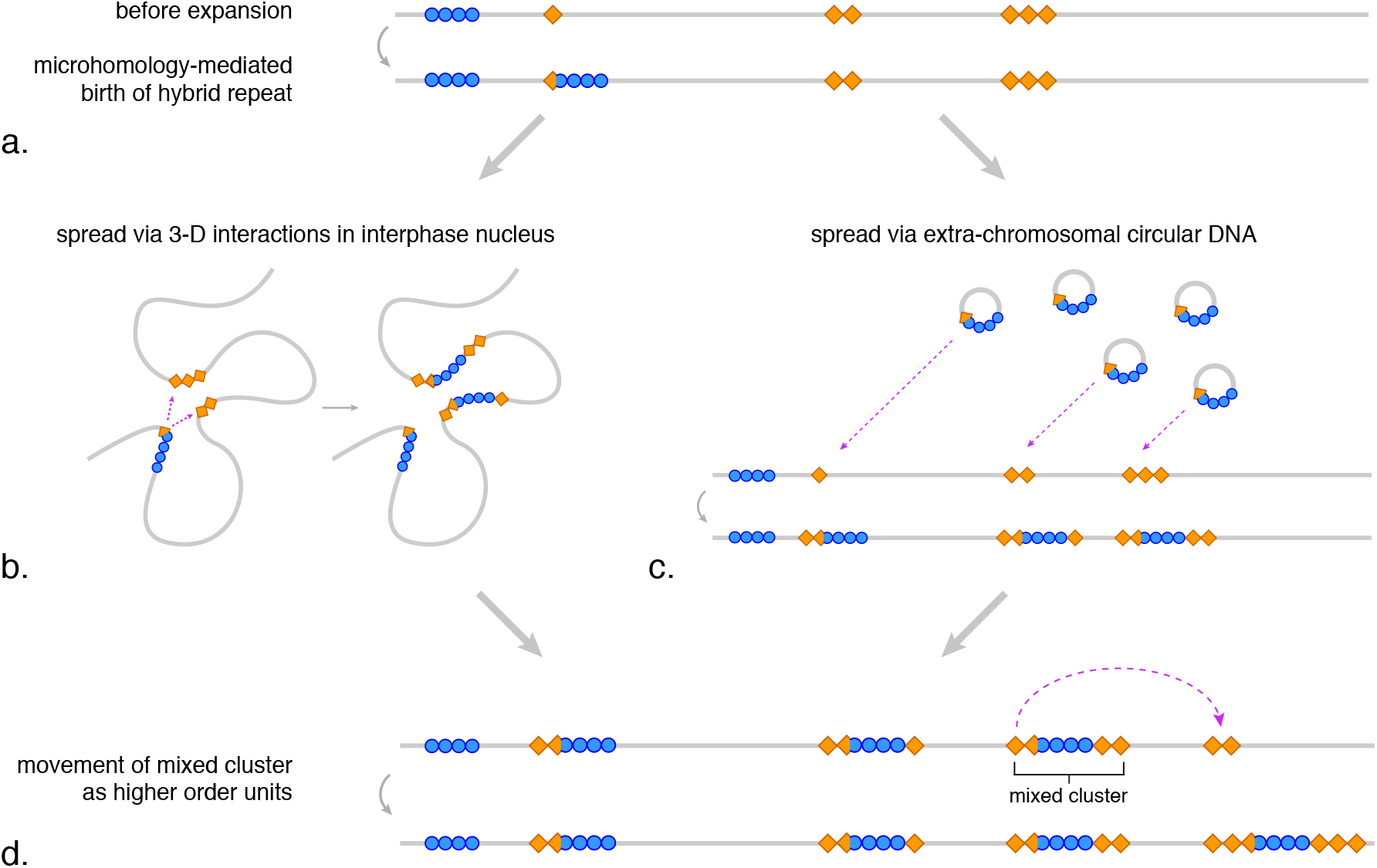
Proposed mechanisms of satDNA dynamics. Blue circles represent an ancestrally rare satellite (*i.e.*, *Rsp-like),* orange diamonds represent an abundant satellite present at many loci (*i.e.*, *1.688*), gray lines represent a fraction of a chromosome that spans many megabases. (a.) illustrates the microhomology-mediated birth of a hybrid repeat formed from the rare+common satellites, facilitating spreading of the rare satellite to loci where the abundant satellite is already present through processes illustrated by b–d. (b.) loci that are physically distant on a linear X chromosome may interact in three-dimensional space within the interphase nucleus, interlocus gene conversion of orange satellite repeats may then facilitate the spread of blue repeats. (c.) satellite DNAs are present on extrachromosomal circular DNAs, which may facilitate their spread to new loci. (d.) after new insertions of the blue satellite, entire mixed clusters may move as higher order units. The mechanisms illustrated in (b) and (c) could also be responsible for the generation of the hybrid repeat (a) and movement of higher order units (d). Not illustrated is the expansion or contraction of a repeat cluster at a given locus due to unequal exchange with a different cluster of the same repeat type.

As described above, the relatively minor *1.688* repeat variants adjacent to the type 1 and type 2 junctions are each shared across multiple *1.688/Rsp-like* clusters (Fig. 5). This suggests either *Rsp-like* has repeatedly inserted into a particular subset of variants in both species, or that the multiple *1.688/Rsp-like* junctions were not formed independently within either species. In the latter scenario, a relatively rare microhomology-mediated event gives rise to a *1.688/Rsp-like* hybrid repeat, which then seeds new *Rsp-like* clusters at loci where *1.688* clusters were already present, facilitated by homology of the *1.688* portion of the novel hybrid repeat. We tested two predictions arising from this model: (1) newly inserted *Rsp-like* clusters would only occur at genomic loci where *1.688* repeats were already present; (2) any *1.688* sequences moving as a higher order repeat along with *Rsp-like* sequences may show discordant phylogenetic relationships with *1.688* repeats already present at the new insertion site.

We tested the above predictions using *D. simulans Rsp-like* clusters with type 1 junctions, focusing on the 12 of 19 clusters that are present at genomic loci where *Rsp-like* clusters are lacking in one or more of the other three study species (*i.e.*, those clusters at cytobands 7-12). We conducted a synteny analysis across species to establish orthology of the 12 clusters. If a *1.688* cluster was present at a syntenic position in the other species, we inferred that *Rsp-like* moved into an existing cluster in *D. simulans.* We found that all 12 new *Rsp-like* clusters in *D. simulans* had *1.688* repeats at that same location in each of the other three species with the exception of a single locus in *D. melanogaster* (Table S3). With the exception of two loci at cytoband 11 in *D. mauritiana,* none of the syntenic loci in the other species have *Rsp-like* repeats (Table S3). The fact that *1.688* clusters were already present at the site of new *Rsp-like* insertions suggests it is sequence homology (and/or microhomology) with *1.688* repeats that is facilitating new insertions. In six of 12 clusters with new insertions, the *1.688* repeat immediately adjacent to the *Rsp-like* junction shows strongly discordant relationship with the other *1.688* repeats in the cluster (Table S3), suggesting that at least a partial *1.688* repeat has moved together with *Rsp-like* repeats.

Our findings from the *1.688/Rsp-like* junction and synteny analyses are consistent with a model in which small regions of microhomology can facilitate the integration of *Rsp-like* into *1.688.* Once this association is created, the rapid spread of *Rsp-like* across the chromosome could be facilitated by hitchhiking with segments of flanking *1.688* repeats (Fig. 6b, c), including through the movement of entire mixed clusters to new locations as a higher-order unit (Fig. 6d).

### Mechanisms underlying spread of clusters to new loci

Two mechanisms that can explain the generation of new clusters as well as the spread of nearly identical repeats: (1) three dimensional interactions in the nucleus creating opportunities for interlocus gene conversion across long linear distances; and (2) the spread of repeats via extrachromosomal circular DNA (eccDNA) to new loci across the X chromosome (Fig. 6). Our reanalysis of *D. melanogaster* Hi-C data (Ogiyama, et al. 2018) provides some evidence of inter-cytoband interactions, particularly across the middle of the X chromosome (*i.e.*, from cytobands 6 through 14) (Fig. S21).

If long-distance gene conversion is facilitated by 3D interactions in the nucleus, we might expect *1.688* repeats and neighboring *Rsp-like* repeats to show a similar pattern of gene conversion. Analysis of sequence similarity of the *1.688* repeats adjacent to these *Rsp-like* clusters showed a mixed pattern, with high sequence similarity among repeats only at cytobands 1, 11, and 12 (Fig. S22). The majority (64.5%) of *1.688* repeats have <95% sequence similarity with any repeat from another cytoband, while the nearest *Rsp-like* repeat shows >95% similarity with repeats from multiple different cytobands. Thus, we find limited evidence of long-distance gene conversion in *1.688* sequences; however, it is possible that the older age and smaller size of *1.688* clusters relative to *Rsp-like* clusters may limit interlocus gene conversion.

### eccDNA as a mechanism of satDNA to new genomic loci

Spread of repeats via eccDNA (extrachromosomal circular DNA) is another (non-mutually exclusive) mechanism that could mediate the spread of *Rsp-like* satellite repeats. We used 2D gel analysis to confirm/show the presence of *1.688* (Cohen, et al. 2003) and *Rsp* eccDNA in *D. melanogaster* (Fig. S23) and then isolated (Fig. S23–24) and sequenced the eccDNA component from all four species. We estimated the abundance of sequences in eccDNA and in the genomic control as reads-per-million (RPM).

We find long-terminal repeats (LTRs) and complex satellites, including *1.688* and *Rsp-like,* are abundant on eccDNAs in all four species (Fig. S25; Fig. 7). In general, we detect a strong correlation between the abundance of a repetitive element in the genome (estimated by RPM for that element in the non-digested genomic DNA control reads) and the abundance of eccDNA reads derived from that repeat. However, some repeats produce more eccDNA than expected given their genomic abundance (Fig. 7). *Rsp-like* repeats are particularly abundant on eccDNA in *D. simulans* (Fig. 7), where they comprise ~3% of the total eccDNA-enriched reads (24.5-fold enrichment over the undigested control), and in *D. sechellia* where they comprise ~4.9% of reads (5.75 enrichment over the undigested control).

**Figure 7.**
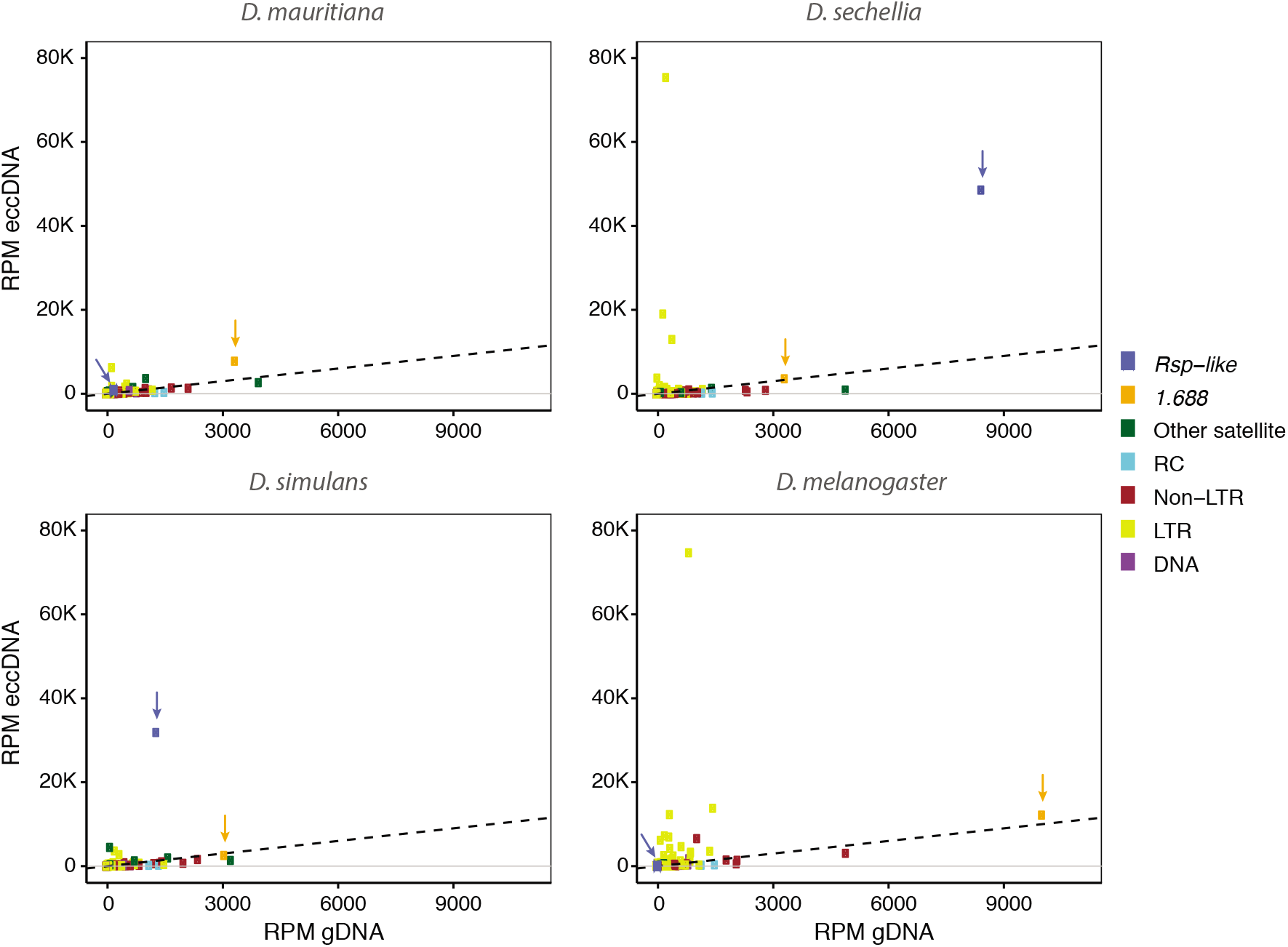
Scatter plot of eccDNA RPM and genomic DNA RPM. Repeats in the genome are categorized into Other satellite (complex satellites except *1.688* and *Rsp-like),* LTR retrotransposon, non-LTR retrotransposon, DNA transposon and rolling-circle (RC) transposon and are shown in different colors. *Rsp-like* (shown in blue) and *1.688* (shown in orange) are indicated by arrows. Dotted lines represent the same abundance of eccDNA and genomic DNA such that dots above the dotted line indicate repeats that are enriched in eccDNA libraries relative to genomic controls.

To determine the genomic source of satellite-derived eccDNAs, we estimated abundance of each sequence variant of *1.688* or *Rsp-like* from euchromatic and heterochromatic loci. We represent the estimated eccDNA abundance on phylogenetic trees by scaling tip labels based on the RPM of each variant (Figs. 3, S7–14). With the exception of *1.688* in *D. sechellia* and *D. mauritiana,* heterochromatic repeat variants produce more eccDNA than euchromatic variants. Consistent with the lack of heterochromatic *Rsp-like* repeats (Larracuente 2014), few eccDNAs map to *D. mauritiana Rsp-like.* Some individual repeats generate more eccDNAs than others, possibly due to sequence composition, chromatin structure, and/or recombination environment. For example, in *D. simulans,* eight euchromatic *Rsp-like* variants from cytoband 5A are enriched for eccDNA (RPM ranges from ~100–600, see light orange tips on Figs. 3, S12). These euchromatic repeats group with the heterochromatic repeats that are also enriched for eccDNA reads (Figs. 3, S12). It is therefore possible that the repeats at 5A may be a result of a recent integration of heterochromatic-derived eccDNA carrying *Rsp-like* repeats.

## DISCUSSION

Our comparative analysis of complex satDNA in high-quality genome assemblies reveals that small blocks of X-linked euchromatic satellites evolve rapidly over short evolutionary time scales. Despite diverging from a common ancestor just 240K years ago (Garrigan, et al. 2012), the *simulans* clade species differ in the total number of repeats, the number of clusters, and in the composition of clusters across syntenic loci (Figs. 1–2, S1; Tables 2, S3). The dynamic evolution of these repeats within the X chromosome euchromatin is similar to the rapid evolution of large blocks of heterochromatic satDNA across whole chromosomes reported in this (Fig. S1), and other studies (Strachan, et al. 1982; Lohe and Brutlag 1987; Lohe and Roberts 1988; Larracuente 2014; Jagannathan, et al. 2017; Wei, et al. 2018). In the euchromatin, however, the expansion, contraction, sequence turnover, and movement of repeats plays out across tens to hundreds of comparatively small loci distributed within a single chromosome. At least some of the differences in repeat abundance between species may be explained by ecology and demographic history. For example, *D. sechellia* is an island endemic with a historically low effective population size (Legrand, et al. 2009) and natural selection may be less efficacious in this species (McBride 2007). Interestingly, this species has larger euchromatic satDNA clusters suggesting that intralocus expansions of repeats may be weakly deleterious, but it does not have more discrete repeat clusters. In contrast to *D. sechellia*, we see the birth of new *Rsp-like* clusters in *D. simulans* and *D. mauritiana* across the X chromosome.

We show that euchromatic satDNAs can proliferate rapidly over short evolutionary timescales. *Rsp-like* repeats recently spread across a ~14 Mb region of the X chromosomes of *D. simulans* and *D. mauritiana,* inserting into existing *1.688* clusters (Figs. 1, 3–4, S3, S8, S10, S12). Although we find that *1.688* has an old history of diversification consistent with previous studies (Waring and Pollack 1987; DiBartolomeis, et al. 1992), our phylogenetic analysis of *1.688* repeats suggests an evolutionary history characterized by long periods of local differentiation among repeats, punctuated by the occasional proliferation of a particular variant, and subsequent local diversification (Figs. 4, S15-16). Thus, our comparative study of repeat patterns in these species reveals satellite proliferation dynamics that may implicate common processes underlying the evolution of both repeat types. These apparent cycles of proliferation and diversification are somewhat analogous to bursts of TE proliferation, except that rather than spreading by encoding proteins to mediate their movement, satDNAs likely spread through recombination-based mechanisms.

### *Mechanisms of* Rsp-like *movement*

We find evidence that microhomology-mediated events generated new hybrid repeats that joined the sequence of a relatively uncommon satellite (*i.e.*, *Rsp-like)* to that of an abundant satellite with a dense distribution across the X chromosome (*i.e*., *1.688).* The birth of new *1.688/Rsp-like* hybrid repeats appears to have occurred independently in *D. simulans* and *D. mauritiana*, and likely multiple times within each species (Figs. 4–5, S17–18, Table S3). Microhomology-mediated repair events are implicated in creating structural rearrangements and chromosomal translocations across organisms (reviewed in McVey and Lee 2008), as well as copy number variations associated with human disease (Hastings, et al. 2009), and gap repair after P-element transpositions in *Drosophila* (Adams, et al. 2003; McVey, et al. 2004). After the initial microhomology-mediated association of the two repeats, the probability of the *Rsp-like* repeats being involved in additional repair events at homologous sequences along the chromosome increased because of their association with *1.688* which is abundant across the X chromosome. Our conclusion that this new association with *1.688* facilitated the spread of *Rsp-like* clusters is supported by both our analysis of junctions and synteny analysis of clusters with new *Rsp-like* insertions (Fig. 5, Table S3). The movement of these higher-order repeats along with intralocus satDNA expansions via unequal exchange further contribute to the fluidity of the repeat landscape (Figs. 5–6). Our investigation of *Rsp-like* proliferation provides a nucleotide-scale illustration of the mechanisms that can account for apparently random, differential amplification of ancestral satellites that leads to species-specific satDNA profiles observed by previous studies *(e.g.,* Mestrovic, et al. 1998; Pons, et al. 2004).

### Mechanisms facilitating long-distance spread of new clusters

Questions remain about the source of the template *Rsp-like* sequences. We discussed two possibilities here: eccDNA reintegration and interlocus gene conversion. Both exploit DNA breaks which out of necessity must be repaired; the nature/timing of the break is an important factor in determining which of the many repair pathways is involved (Scully, et al. 2019). The complexity of the sequences observed in the *Rsp-like/1.688* variable junctions could implicate pathways such as FoSteS (fork stalling and template switching, Lee, et al. 2007), or MMBIR (microhomology-mediated break-induced replication, Hastings, et al. 2009). Both of these repair pathways occur during aberrant DNA replication and can involve multiple template switches facilitated by microhomology. Alternatively, during double-strand break (DSB) repair, synthesis-dependent strand annealing with an interlocus template switch may result in gene conversion events *(e.g.,* Smith, et al. 2007) that insert *Rsp-like* sequences into existing *1.688* clusters. Similar events occur at the yeast MAT locus during gene conversion, where interchromosomal template switches occur even between divergent sequences, and these events can proceed based on microhomologies as small as 2 bp (Tsaponina and Haber 2014). DNA prone to forming secondary structures *(e.g.,* non-B form DNA like hairpins or G quartets) can cause replication fork collapse that leads to DSB formation (reviewed in Mirkin and Mirkin 2007). Blocks of complex satDNAs may be enriched for sequences that form secondary structures and therefore may have elevated rates of DSBs compared to single-copy sequences. Elevated rates of DSB may make it more likely to observe non-homologous recombination-mediated repair events resulting in complex rearrangements, differences in repeat copy number and, as we describe here, the colonization of repeats at new genomic positions across large physical distances.

We show that complex satellites are abundant on eccDNA (Figs. 7, S23–25), and map eccDNA reads to the specific repeat variants from which these circles arise (Figs. 3, S7–14). While the abundance of most eccDNAs correlates with their genomic abundance, some repeats, such as *Rsp-like* in *D. simulans,* generate a disproportionate amount of eccDNAs. The formation of eccDNA may depend on DNA sequence, organization *(e.g.,* repetitive versus unique), chromatin status, and possibly its higher order structure. It is possible that the high abundance of *Rsp-like* derived eccDNA suggests that this satellite is unstable at the chromatin level, or more prone to DSB. EccDNA formation exploits different methods of DNA damage repair, including HR (using solo LTRs, (Gresham, et al. 2010), MMEJ (Shibata, et al. 2012; Moller, et al. 2015), and NHEJ (van Loon, et al. 1994). The repetitive nature of *1.688* and *Rsp-like* makes it difficult to examine junctions in the extrachromosomal circles themselves. We do find evidence suggesting that HR can give rise to *Rsp-like* circles, however. An eccDNA arising from an intrachromatid exchange event between repeats within the same array, followed by the reintegration of that eccDNA at a new genomic location, could generate new arrays where the first and last repeat are truncated, but together would form a complete monomer. We see this pattern in four of the new *Rsp-like* arrays in *D. simulans* (Dsimpre1A-a, Dsimpre1A-b, Dsimpre1A-c, Dsim1A-1; Fig. 5) and two arrays in *D. mauritiana* (Dmau1A-4, Dmau1A-6; Fig. 5). It is thus conceivable that eccDNAs are involved in the generation of new *Rsp-like* clusters. Our finding that satDNAs and LTRs are enriched on circles is consistent with other studies showing that repeats generate eccDNA (Cohen, et al. 2003; Cohen, et al. 2006; Navratilova, et al. 2008; Cohen and Segal 2009; Moller, et al. 2015; Lanciano, et al. 2017; Shoura, et al. 2017). EccDNAs may be a source of genomic plasticity within species (Gaubatz 1990); we suspect that they also played a role in the proliferation of satDNAs in the *simulans* clade, thus contributing to X-linked repeat divergence between these species. Experimental approaches will help explicitly test the hypothesis that satDNA-derived eccDNAs reintegrate in the genome.

Interactions in the 3D nucleus may also contribute to movement of satDNA by facilitating interlocus gene conversion events between loci far apart on a linear chromosome, including through heterochromatin/euchromatin interactions (Lee, et al. 2019). Although our data are not suited to directly test this hypothesis, we find indirect evidence that long-distance interactions may occur across the X euchromatin through reanalysis of *D. melanogaster* Hi-C data and by searching for signatures of recent gene conversion in *1.688* repeats flanking regions with new *Rsp-like* insertions (Fig. S21–S22). If these long-distance interactions in the 3D nucleus are conserved between species, this may account for the similar but independent spread of satDNAs to distant loci that we see in *D. simulans* and *D. mauritiana*. Data on long-range 3D chromosome interactions *(e.g.,* Hi-C) in the *simulans* clade species will be important for testing this hypothesis and for understanding the role of interlocus gene conversion in satDNA movement.

### Functional consequences of rapid satDNA evolution

A growing body of research suggests that shifts in satellite abundance and location may have consequences for genome evolution. Large scale rearrangements or divergence in heterochromatic satDNA may lead to hybrid incompatibilities. In *D. melanogaster* a heterochromatic block of *1.688* satDNA is associated with embryonic lethality in *D. melanogaster – D. simulans* hybrids (Ferree and Barbash 2009; Ferree and Prasad 2012) through mechanisms that we do not yet understand. However, even variation in small euchromatic satDNAs can have measurable effects on gene regulation and thus may be important for genome evolution. Short tandem repeats in vertebrate genomes can affect gene regulation by acting as binding sites for transcription factors (Rockman and Wray 2002; Gemayel, et al. 2010). Additionally, repeats can have an impact on local chromatin, which may affect nearby gene expression *(e.g.,* Feliciello, et al. 2015). Novel TE insertions can cause small RNA-mediated changes in chromatin *(e.g.,* H3K9me2) that can spread to nearby regions and alter local gene expression (Lee and Karpen 2017). In *D. melanogaster,* siRNA mediated chromatin modifications at some *1.688* repeats play a role in X chromosome recognition during dosage compensation (Menon, et al. 2014; Joshi and Meller 2017; Deshpande and Meller 2018). The turnover in repeat composition in *D. simulans* and *D. mauritiana* at loci *(e.g.,* Fig. 2) with demonstrated effects on chromatin and MSL recruitment (Joshi and Meller 2017; Deshpande and Meller 2018) raises the possibility that dynamic evolution of euchromatic satDNAs may have functional consequences for dosage compensation.

Understanding of the molecular mechanisms that drive rapid expansion, movement, and rearrangement of satDNAs across the genome is a necessary step in determining the functional and evolutionary consequences of rapid satDNA evolution. In addition to fine-scale mapping of satDNA evolution in a comparative framework, we present initial insights as to the mechanisms that shape the proliferation and movement over short time scales. Future work that includes population data will be important for disentangling species vs population-level variation and addressing whether natural selection plays a role in satDNA evolution within and across loci. We suspect that the rapid satDNA dynamics in one genome compartment *(e.g.,* heterochromatin) may drive corresponding changes in the other genome compartment *(e.g.,* X-linked euchromatin).

Future work on the evolutionary forces driving rapid satDNA evolution *(e.g.,* molecular drive (Dover 1982), meiotic drive (Henikoff, et al. 2001)), and the molecular and physical interactions between heterochromatin and euchromatin *(e.g.,* Lee, et al. 2019), will help reveal the broad consequences for rapid satDNA evolution.

## MATERIALS AND METHODS

### Repeat annotation

Repeat annotations were performed as described in (Chakraborty, et al., unpublished). Briefly, we constructed a custom repeat library by downloading the latest repetitive element release for *Drosophila* from RepBase and added custom satellite annotations. We manually checked our library for redundancies and miscategorizations. We used our custom library with RepeatMasker version 4.0.5 using permissive parameters to annotate the assemblies. We merged our repeat annotations with gene annotations constructed in Maker version 2.31.9 (for the *simulans* clade species) (Cantarel, et al. 2008) or downloaded from Flybase (for *D. melanogaster)* (Thurmond, et al. 2019).

We used custom Perl scripts to define clusters of satellites on the X chromosome and to determine the closest neighboring annotations. We defined clusters as two or more monomers of a given satellite within 500 bp of each other, though some analyses we also included single monomers. We grouped clusters according to cytoband (FlyBase annotation v6.03; ftp://ftp.flybase.net/releases/FB2014_06/precomputed_files/map_conversion/). We used custom scripts to translate the coordinates of cytoband boundaries from *Drosophila melanogaster* to the other three species with the following workflow. We extracted 30K bases adjacent to the coordinate of each cytoband sub-division in the *D. melanogaster* assembly and used that sequence as a query in a BLAST search against repeat-masked versions of the *simulans* clade species genomes. To obtain rough boundaries of *D. melanogaster* cytobands in each *simulans* clade species, we defined the proximal-most boundary as the proximal coordinate of the first hit (>1 kb in length) from each cytoband region. We defined the distal boundary arbitrarily as one base less than the proximal coordinate of the next cytoband.

### *Analysis of* 1.688/Rsp-like *junctions*

We tested the hypothesis that short regions of microhomology could facilitate the insertion of *Rsp-like* repeats at new genomic loci using two complementary approaches: (1) through extensive visual examination of *1.688/Rsp-like* junctions in *D. simulans* and *D. mauritiana* in the context of multi-sequence alignments as well as the X chromosome assembly in Geneious v8.1.6. (2) we used MEME (Bailey, et al. 2015) to computationally detect motifs that are enriched at the edges of new *Rsp-like* clusters. Additional details are provided in Supplemental Material.

### *Analysis of syntenic* 1.688 *clusters with* Rsp-like *insertions in* D. simulans

We tested the prediction that new *Rsp-like* clusters would insert only at loci where *1.688* clusters were already present by extracting 5 kb of sequence immediately upstream and downstream of the loci containing a mixed *1.688/Rsp-like* cluster in *D. simulans.* We determined the orthologous position of these flanking sequences in the other three study species by using the flanks as BLAST query sequences which we searched against custom BLAST databases built from the assemblies of the other species. We accepted best hits as orthologous sequences only if they were reciprocal best hits when BLASTed back against the *D. simulans* genome assembly. We then navigated to the orthologous flanking sequences of each cluster to determine whether a *1.688* cluster was present at that locus in the three other study species.

We tested for discordant phylogenetic relationships among *1.688* repeats in clusters with new *Rsp-like* insertions in *D. simulans* by extracting *1.688* repeats surrounding the *Rsp-like* insertion and flagging those sequences in a phylogenetic analysis in which they were included with all *1.688* euchromatic repeats from *D. simulans.* We extracted flanking sequences, generated custom BLAST databases, conducted BLAST searches, and extracted relevant *1.688* monomers in Geneious v.8.1.9. For both of the above tests, we used as models those *Rsp-like* clusters that show the dominant junction signature in *D. simulans* (Fig. 5), with a focus on 12 clusters that are present at genomic loci where *Rsp-like* clusters are lacking in one or more of the other three study species (*i.e.*, those clusters at cytobands 7-12).

### Extrachromosomal circular DNA isolation and sequencing

Genomic DNA was isolated from 20 five-day adult females (20-25 mg) from *D. melanogaster* (strain iso 1), *D. mauritiana*, (strain 12), *D. sechellia* (strain C), and *D. simulans* (strain XD1) using standard phenol-chloroform extractions. The DNAs were ethanol precipitated and resuspended in 10 mM Tris-EDTA, pH 8.0. The concentrations were determined by Qubit fluorometric quantification. 200 ng of each genomic DNA was subjected to exoV (New England Biolabs) digestion as described by (Shoura, et al. 2017). In short, after digestion at 37° for 24 hours, the DNAs were incubated at 70° for 30 minutes. Additional buffer, ATP, and exoV were then added and the samples incubated at 37° for another 24 hours. The process was repeated for a total of 4, 24 hour incubations with exoV. The concentration of the remaining DNA was determined by Qubit. Following circle isolation, we prepared libraries of circle-enriched and whole genomic control samples using NEBNext FS DNA Ultra II Library Prep Kit (New England Biolabs) using protocol modifications outlined in (Sproul and Maddison 2017). Libraries were pooled and sequenced on the same 150-base paired-end lane of an Illumina HiSeq 4000 by GENEWIZ laboratories (South Plainfield, NJ, USA). Additional methods for library preparation and mapping variants of eccDNA to phylogenetic trees is provided in Supplemental Material.

## ACKNOWLEDGEMENTS

We thank Massa Shoura (Stanford University) for advice on eccDNA isolation. We also thank Tom Eickbush, Jack Werren, and Ching-Ho Chang for helpful feedback on the study. Illumina genomic DNA and eccDNA raw reads for each species will be available in NCBI’s SRA (accession is forthcoming). All data files and code for analysis and producing plots are deposited in GitHub (https://github.com/LarracuenteLab/simulans_clade_satDNA_evolution) and in the Dryad Digital Repository (doi is forthcoming). This work was supported by National Institutes of Health General Medical Sciences grant (R35GM119515 to AML), the University of Rochester, and a Stephen Biggar and Elisabeth Asaro Fellowship in Data Science. JSS is supported by an NSF Postdoctoral Research Fellowship in Biology (DBI-1811930).

