## Supplement for "Dynamic evolution of euchromatic satellites on the X chromosome in *Drosophila melanogaster* and the *simulans* clade"

### SUPPLEMENTARY METHODS

*Fluorescence in-situ hybridization*

We studied broad-scale dynamics of complex satellites by mapping the location of *1.688* and *Rsp-like* repeats on *Drosophila* chromosomes using FISH protocols outlined in Larracuente and Ferree (Larracuente and Ferree 2015). Briefly, larval brains were dissected in 1× PBS, treated with a hypotonic solution (0.5% sodium citrate) and fixed in 1.8% paraformaldehyde, 45% acetic acid, and dehydrated in ethanol. For salivary glands, the same procedure was followed except for the treatment with hypotonic solution. Additional We generated biotin- and digoxigenin-labeled probes using nick translation on gel-extracted PCR products from 360-bp (*D. simulans* DNA; 360F:'ACTCCTTCTTGCTCTCTGACCA'; 360R:'CATTTTGTACTCCTTACAACCAATACTA') (Ferree and Barbash 2009), and *Rsp-like* (*D. sechellia* DNA; Rsp-likeF:'ACTGATTATCATCGCCTGGT'; Rsp-likeR:'GTAACTCCAGTTCGCCTGGT) (Larracuente 2014). For the *D. melanogaster 1.688* probe, we generated biotin-labeled probes using nick translation on gel-extracted PCR products from 260-bp repeats ( 260F: 5′-TGGAAATTTAATTACGAGCT-3′; 260R: 5′-ATGAAACTGTGTTCAACAAT-3′) (Abad, et al. 2000), which cross hybridize with all heterochromatic 1.688 repeats (Khost, et al. 2017). We made the *simulans* clade *1.688* probe in the same way (360F 5'-ACTCCTTCTTGCTCTCTGACCA-3', 360R 5'-CATTTTGTACTCCTTACAACCAATACTA-3' (Ferree and Barbash 2009)). Probes were hybridized overnight at 30°C, washed in 4× SSCT and 0.1×SSC, blocked in a BSA solution, and treated with 1:100 Rhodamine-avadin (Roche) and 1:100 anti-dig fluorescein (Roche), with final washes in 4× SSCT and 0.1× SSC. Slides were mounted in Vecta-Shield with DAPI (Vector Laboratories), visualized on a Leica DM5500 upright fluorescence microscope at 100×, imaged with a Hamamatsu Orca R2 CCD camera, and analyzed using Leica's LAX software.

*Evolutionary relationship of satDNAs within and among species*

We compared evolutionary histories of *1.688* and *Rsp-like* by generating phylogenetic trees for each repeat within species (referred to in the text as 'within-species' trees). We compared patterns across the resulting trees by focusing on four aspects of the topologies: (1) general patterns of nodal support and branch lengths; (2) the relationship of repeats in euchromatic vs heterochromatic genome regions; (3) the fraction of highly-supported clades for which all descendants are repeats from the same cytoband, or two adjacent cytobands; this comparison is intended to test the null hypothesis that repeats from physically nearby location (*e.g*., those within a cluster, or from nearby clusters) are expected to homogenize via gene conversion and show greater sequence similarity than physically distant clusters – deviation from this null model (*i.e.,* a tree with a low fraction of repeats showing local homogenization) could indicate recent spread of repeats to new loci where local homogenization and subsequent differentiation from other clusters has not yet had time to accumulate; (4) the presence of clades containing repeats from cytobands that are physically distant (*i.e*., non-adjacent) relative to the linear organization of the X chromosome, which could indicate historical exchange events that span large physical distances relative to the linear organization of the X chromosome. In addition to the above within-species trees, we generated 'all-species' trees for both *1.688* and *Rsp-like* repeats, which combined monomers from all four species in the same analysis. The all-species trees allowed additional insight as to the relative timing of diversification within each satellite type, and to test whether interlocus expansions of *Rsp-like* repeats in the *simulans* clade species share a common origin, or occurred independently.

For all phylogenetic analyses, we aligned repeats using MAFFT (Katoh and Standley 2014) in Geneious v8.1.6 with the "auto" option, which selects the most efficient algorithm based on the number of input sequences. Prior to alignment, we filtered the data to exclude monomers originating in small clusters (*i.e.*, those with ≤ two monomers) and monomers below a minimum length (*i.e.*, ≤ 100bp for *Rsp-like*, ≤300bp for *1.688*). As outgroup sequences, we used consensus sequences of *Rsp-like* and *1.688* repeats from *Drosophila erecta*, a closely related species that has a sister relationship to all study taxa. We used RAxML v8.2.11 to infer maximum likelihood trees with GTR+gamma as the model of evolution, and conducted bootstrap analysis using the --autoMRE option to automatically determine the optimal number of bootstrap replicates (Stamatakis 2014) which is recommended for large data sets in the program documentation. The resulting trees were plotted and stylized using APE and ggtree in R (Yu, et al. 2018) and Adobe Illustrator.

*Cluster age estimation*

We analyzed differential patterns of sequence divergence to infer patterns of gene conversion within a repeat cluster and estimate the relative age of a given cluster. This analysis was designed to test our conclusion that *Rsp-like* clusters in *D. simulans* and *D. mauritiana* at novel loci are due to new insertions against the alternative that these are actually older clusters that were lost in the other species, which are being maintained homogeneous by long-distance gene conversion. We reasoned that gene conversion (and unequal exchange) is more likely to happen in the center of repeat clusters leading to the homogenization of repeats in the cluster center and an accumulation of sequence variants towards the edges of the cluster, in accordance with the accretion model of repeat evolution (McAllister and Werren 1999). We thus expect older clusters to have low sequence divergence among repeats within the center of the cluster and high divergence between the first/last repeats and the center of the cluster. A "new" cluster would not have had time to accrue mutations and/or homogenize its center repeats through gene conversion, and therefore we would not expect to increased sequence divergence in repeats at the edge of the cluster relative to those in the middle.

We calculated a pairwise distance matrix from the alignment of repeats within each cluster. For each cluster with at least 4 repeats, we divided the cluster into three segments: first (first complete monomer), last (last complete monomer), and center (all repeats in between first and last). We then compared the genetic distance between the first/last repeats and the center vs the average genetic distance among repeats within the center. We define a metric, dY, as the maximum of two comparisons: (1) the first-center distance vs the within-center distance; and (2) the last-center distance vs within-center distance. The plots of first-center vs center-center vs last-center average distances were visualized as scatterplots using R (Fig S20). We then used the difference between first/last-center and center-center distances to partition clusters into old or new: new clusters should have small differences between first/last and center repeats, while older clusters should have larger differences. Based on the distribution of dY, we use $dY<0.05$ as the threshold below which clusters are inferred to be "young". Changing the dY threshold for "new" does not qualitatively change the results and interpretations. Clusters that we can confidently infer to be shared between the *simulans* clade species (e.g. the *Rsp-like* cluster at 4B) and are thus "old" have a high dY, which supports the validity of our approach.

To prepare the data for cluster age estimation, we extracted repeat monomers from the assemblies of *D. melanogaster* (Khost, et al. 2017) and each species of the *simulans* clade (Chakraborty, et al.). We aligned repeats for each cluster using ClustalO (Sievers and Higgins 2014) and MUSCLE (Edgar 2004), followed by manual curation of alignments to correct for mis-alignment or mis-annotations in Geneious v8.1.6. We generated distance matrices from alignments of each cluster with the dist.dna function in APE (Paradis and Schliep 2019) using the F81 model of sequence evolution (Felsenstein 1981). We chose the model of evolution following the results of model selection analysis conducted for all cluster alignments using ModelTest (Posada and Crandall 1998) as implemented in the R package Phangorn (Schliep 2011), which identified the F81 model as the best fitting model for the majority of clusters under the Bayesian information criterion.

*Junction analysis with MEME*

We used MEME v4.10.0 (Bailey, et al. 2015) to identify sequence motifs near the boundaries of *Rsp-like* satellite clusters in X chromosome euchromatin for the *simulans* clade and *D. melanogaster*. We generated the MEME dataset by extracting 120 bases on either side of all *Rsp-like* clusters such that 20 bases overlapped the *Rsp-like* annotation, and 100 bases extended into the sequence flanking the annotation. We analyzed sequences in MEME to search for the top 25 motifs ranging from 8 to 20 bases. We mapped motifs identified by MEME to the X chromosome of each species using FIMO v4.10.0 (Grant, et al. 2011). We determined which genomic attributes overlapped motif coordinates by comparing all motif coordinates generated by FIMO to the gff file containing X chromosome attribute annotations using custom scripts written in bash and R. We identified genomic attributes that were enriched for each motif relative to the rest of the X chromosome by calculating the total fraction of the X chromosome covered by a given attribute and then calculating expected coverage of each motif in that attribute given the total coverage of that motif on the X chromosome. We compared this expected value to the observed value obtained from analysis of FIMO results described above using a Fisher's exact test (using the 'fisher.test' function) followed by a Bonferroni correction for multiple comparisons (using the 'p.adjust' function) in R. We visualized motifs relative to sequence alignments of *Rsp-like* and *1.688* clusters in Geneious v8.1.6.

*Hi-C analysis of 3D interactions in D. melanogaster embryo*

We used a publicly available Hi-C dataset from stage 16 embryos (Gene Expression Omnibus accession number GSE103625) to test the 3D interactions among satellite repeats in *D. melanogaster* (Ogiyama, et al. 2018). We mapped Hi-C raw sequence reads to the r6 reference genome, and processed the output with the HiC-Pro pipeline (Servant, et al. 2015) to obtain contact matrix at 10kb resolution (default parameters). We summarized results from the contact matrix in R using the Biocircos v0.3.4 (Chen and Cui 2019). We plotted inter-cytoband interactions using a cutoff of normalized interaction counts > 40 in 10-kb windows and excluded the *1.688* sequences themselves to avoid potential mappability issues.

*Testing for gene conversion at* 1.688/Rsp-like *junctions*

To test whether *1.688* clusters near *Rsp-like* clusters show evidence of recent gene conversion, we created all-by-all distance matrices of *Rsp-like* repeats. In addition, we created a similar distance matrix of all *1.688* repeats that are within 100 bases of a *Rsp-like* cluster. We plotted each distance matrix as a circular plot (similar to genome synteny plots) using BioCircos v0.3.4 (Chen and Cui 2019). In the resulting plot, each repeat is grouped by cytoband, and any repeats with genetic distances $\leq 0.05$ have connecting lines drawn between their positions on the circle. Both *1.688* and *Rsp-like* plots were made on the same cytoband scale. This allowed us to overlay the *Rsp-like* and *1.688* plots in order to compare their patterns of sequence divergence at adjacent positions. We only plotted clusters with more than two repeats.

*Verification of eccDNA in exoV digestion*

Aliquots of the undigested genomic DNAs were diluted to the comparable volumes of samples after exoV digestion. A dilution series was then made for PCR analysis of both the exoV digested and the undigested DNAs. Primers used included those for rp49 [5'-CAGCATACAGGCCCAAGATC-3', 5'-CAGTAAACGCGGTTCTGCATG-3'], tRNA(lysine) [5'-CTAGCTCAGTCGGTAGAGCATGA-3', 5'-CCAACGTGGGGCTCGAAC -3'], mitochondria COXI [5'-GATCAAACAAATAAAGGTATACG-3', 5'-GTTCCATGTAAAGTAGCTAATC-3'], 5S [5'-GCCAACGACCATACCACG-3', 5'-GTGGACGAGGCCAACAAC-3'], *Rsp* [5'-GGAAAATCACCCATTTTGATCGC-3', 5'-CCGAATTCAAGTACCAGAC-3'], *Rsp-like* [5'-ACTGATTATCATCGCCTGGT-3', 5'-GTAACTCCAGTTCGCCTGGT-3'], *1.688* [mel 5'-5'GTTTTGAGCAGCTAATTACC-3', mel 5'TATTCTTACATCTATGTGACC-3' (Usakin, et al. 2007) and sech 5'-ACTCCTTCTTGCTCTCTGACCA-3', sech 5'-CATTTTGTACTCCTTACAACCAATACTA-3'].

*2D gel analysis*

Genomic DNA was isolated from the Raleigh 370 strain of *D. melanogaster* as described above. 10 ug of DNA was fractionated by electrophoresis as described by Cohen, et al. (2003).The DNA was then depurinated, denatured, and neutralized before being transferred overnight in high salt (20 X SSC/ 1 M NH4Acetate) to a nylon membrane (Biodyne, ThermoScientific). DNA was UV crosslinked and hybridizations were done overnight at 55°C in North2South hybridization buffer (ThermoScientific). Biotinylated RNA probes were generated from *Rsp* or *1.688* PCR generated amplicons as described previously (Khost, et al. 2017). The hybridized membrane was processed as recommended for the Chemiluminescent Nucleic Acid Detection Module (ThermoScientific), and the signal recorded on a ChemiDoc XR+ (BioRad).

*eccDNA library preparation and sequencing*

We prepared eccDNA-enriched samples and genomic DNA control samples for Illumina sequencing using a NEBNext FS DNA Ultra II Library Prep Kit (New England Biolabs). To control for bias associated with differential PCR amplification among libraries, we used results from an initial round of library preparation to understand variation in library yield between eccDNA isolates and control samples. Initial bioanalysis traces revealed over-amplification in our genomic controls and probable primer/adapter dimers in our eccDNA-enriched samples. To eliminate over-amplification, we halved the amount of input in our control samples and used protocol modifications outlined in (Sproul and Maddison 2017) to reduce adapter dimer content and maximize yield of eccDNA-enriched samples. We generated final libraries using 2 ng of input for eccDNA-enriched samples and 1 ng of input for control samples, with 13 amplification cycles for all samples to minimize amplification bias and allow comparison between samples. Bioanalysis of resulting libraries showed clean traces for all samples (*e.g.*, no evidence of primer/adapter dimer peaks or over amplification).

Libraries were pooled and sequenced on the same 150-base paired-end lane of an Illumina HiSeq 4000 by GENEWIZ laboratories (South Plainfield, NJ, USA). Reads from the control and enriched samples were evaluated using FastQC and trimmed using Trimgalore, then were mapped to the genome using Bowtie2 default parameters. For the repeat composition analysis (Figs. 7 and S24), we used a heterochromatin enriched assembly for *D. melanogaster* (Chang and Larracuente 2019), which has more complete repeat information in heterochromatin regions. Based on our repeat annotations, we calculated the reads per million (RPM) for each repeat using a custom python script. We calculated relative abundance of eccDNA for each repeat in each species by normalizing to its own undigested genomic DNA control. We excluded simple tandem satellite repeats (monomers of 5-12 bp) following analysis because of Illumina read bias from library preparation. To estimate the linear DNA contamination in our eccDNA enriched library, we calculated the RPM values for all genes in the genome (excluding histone cluster and rDNA loci) using HTSeq-count (Anders, et al. 2015), and we found that the mean and median of gene RPMs in eccDNA enriched libraries are ~5%~20% of that in undigested genomic DNA control libraries for all species, suggesting effective enrichment of eccDNA in our eccDNA libraries.

#### SUPPLEMENTARY RESULTS

##### *Phylogenetic results*

Examination of within-species and all-species phylogenetic trees of satellite repeats led to four major findings. (1) Heterochromatic repeats form clades that are generally separate from euchromatic repeats for both satellites in all species except *D. sechellia*, for which euchromatic and heterochromatic repeats are interspersed in both *1.688* and *Rsp-like* (Figs. S7–14). (2) *D. sechellia* and *D. mauritiana* (especially the former) show repeated evidence of intralocus expansion of repeats (Figs. S15–16). (3) *1.688* euchromatic repeats have a relatively old diversification history that largely pre-dates the speciation events that gave rise to the study species (Figs. 3–4, S7, S9, S11, S13, S15–16). This contrasts with *Rsp-like*, which shows evidence of relatively recent diversification, particularly in the *simulans* clade species (Figs. 3–4, S8, S10, S12, S14, S17–18). (4) *Rsp-like* repeats show evidence of two major expansions (Figs. 3–4, S8, S10, S12, S17–18), which encompass large physical distances across the X chromosome (*i.e.*, 'interlocus' expansions) and mainly occurred independently in *D. simulans* and *D. mauritiana*. The latter two findings are discussed in additional detail in the next two paragraphs.

Comparison of within-species trees for *Rsp-like* and *1.688* show contrasting patterns of branch length, nodal support, and local differentiation of repeats, which all indicate an older history of *1.688* diversification (*i.e.*, finding three, above; Figs. 3–4, S7–13; Table S1). The *1.688* all-species tree also supports this conclusion as it reveals deeply divergent clades separating extant *1.688* variants (Fig. 4). Several major *1.688* clades contain repeats from cytobands spanning large physical distances across the X (*e.g.*, the basal clade contains repeats from cytobands 1, 3,

9, and 11 from all four study species; Figs. S15–16). Together they suggest a recurrent history in which an ancestral variant proliferated, spread across the X chromosome, and subsequently underwent local diversification. This diversification of *1.688* repeats largely pre-dated the speciation events that gave rise to the four species (Figs. 4, S15–16). We reach this conclusion upon finding repeated instances of cytoband-specific, well-supported clades comprised of repeats from all four species, with a branching pattern that matches the evolutionary history of the species (*i.e.*, *D. melanogaster* repeats forming a clade sister to the repeats of the *simulans* clade species; Figs. S15–S16). The relative ages of *1.688* and *Rsp-like* repeats are further supported by our estimates of cluster age based on within-cluster repeat divergence (Figs. S19–S20, Table S2). Finally, our observation that *1.688* is a relatively old satellite is consistent with similar conclusions from previous studies (Hsieh and Brutlag 1979; Waring and Pollack 1987; DiBartolomeis, et al. 1992).

The *Rsp-like* all-species tree shows evidence of two major interlocus expansions of *Rsp-like* repeats (*i.e.*, finding four, above) which occur as clades containing hundreds of repeats separated by short branches. One interlocus expansion occurred in the ancestor of the *simulans* clade (Figs. 4 and S17); the second occurred within *D. simulans* alone (Figs. 4 and S18). Repeats from *D. mauritiana* account for 58.9% (n=178) of terminals in the sim-clade expansion and include repeats from cytoband 1, 3, 5, 11, and 12. Repeats from cytobands 1 and 2 in *D. sechellia* make up 25.8% (n=78) of terminals in the sim-clade expansion. The remaining 15.2% (n=46) of repeats are from cytobands 2–5 in *D. simulans*. The sim-specific expansion comprises 226 *D. simulans* repeats from cytobands 1, 3, 7, 8, 9, 10, 11, and 12, all separated by extremely short branches (sim-specific expansion; Figs. 4 and S17); this expansion accounts for most of the increased *Rsp-like* repeats in *D. simulans* relative to all other species. Although the effects of gene conversion could have erased evidence of a common origin of the expansions, the patterns in the all-species tree (Figs. 4 and S17–18) suggest that the *Rsp-like* repeats proximal to cytoband 6 in *D. simulans* and *D. mauritiana* arose through independent interlocus expansions.

*Comparison of within-region patterns of gene conversion*

In all four species, 48.8–78.7% of euchromatic *1.688* repeats form highly-supported clades with other repeats from the same, or an immediately adjacent cytoband (Table S1). This signature suggests that a large fraction of *1.688* repeats show local homogenization through gene conversion and evolve separately from physically distant repeats. We see a different pattern in *Rsp-like* trees for all species, which showed only 2.4–31.8% of euchromatic repeats in highly supported clades with other nearby repeats. The difference was most notable in *D. simulans, D. mauritiana*, and *D. sechellia* (2.4%, 10.3%, and 14.3% of repeats respectively) (Table S1). In the three *simulans* clade species the regions in the *Rsp-like* trees are characterized by short branches with low nodal support and correspond to the *Rsp-like* expansions noted in the all-species tree (Figs. 3, 4, and S8, S10, S12, S14). This pattern of a reduced signature of local gene conversion in *Rsp-like* repeats against a backdrop where *1.688* repeats show higher rates of gene conversion is consistent with the conclusion that *Rsp-like* has a recent history of movement across the X chromosome and clusters have not yet had time to accumulate locus-specific differences.

Rsp-like *clusters are younger than* 1.688 *clusters*

Cluster age estimates show that on average euchromatic *Rsp-like* clusters are younger than *1.688* clusters, particularly in *D. simulans* (17/25 clusters have dY<0.05) and *D. mauritiana* (11/17 clusters have dY<0.05) (Fig. S19; Table S2). In particular, the *Rsp-like* clusters in *D. simulans* and *D. mauritiana* at genomic regions lacking such clusters in *D. sechellia* and *D. melanogaster* have low dY scores. This finding is consistent with phylogenetic evidence that *Rsp-like* repeats have diversified in the recent evolutionary history of the *simulans* clade and supports the hypothesis that the presence of proximal *Rsp-like* clusters in *D. simulans* and *D. mauritiana* are due to new insertions rather than long-distance gene conversion of ancestral repeats that have been recently lost *in D. sechellia* and *D. melanogaster*. We note that a potential source of bias in our cluster age estimate analysis could arise if clusters being measured are actually higher order repeats (e.g., dimers or trimers) in which individual monomers are divergent from each other. Our approach would over-estimate the relative age of such clusters.

*Analysis of* 1.688/Rsp-like *junctions*

In addition to visual examination of junctions described in the main text we identified motifs that are enriched near junctions using MEME (Bailey, et al. 2015) and FIMO (Grant, et al. 2011). This analysis showed that motifs that are abundant at *1.688/Rsp-like* junctions are notably enriched in loci annotated as *1.688* repeats, *Rsp-like* repeats, and genes (Fig. S26). In addition, MEME results were consistent with the results of our visual inspection of junctions in that one of the abundant motifs identified containing the 'TGGTACC' microhomology was present at junction sequences.

*Testing for long-distance interaction in D. melanogaster with Hi-C*

In *D. melanogaster* (Ogiyama, et al. 2018), we explored Hi-C data to make a more direct test of long-distance interactions. We found weak evidence of inter-cytoband interactions, especially across the middle of the X (i.e., from cytobands 6 through 14) (Fig. S21). We plotted inter-band interactions using a cutoff of normalized interaction counts > 40 in 10-kb windows. The mean normalized interaction scores across the whole genome and across the X chromosome are 6 and 9, respectively. A cutoff of 40 corresponds to the 98.8th and $97.7^{th}$ percentile of interaction scores for the whole genome and the X, respectively. Although part of this pipeline is to only use uniquely mapped reads, we excluded *1.688* sequences from these interactions analyses to avoid potential mappability issues with repeats.

#### SUPPLEMENTARY FIGURE CAPTIONS

**Figure S1. Complex satellites in the heterochromatin of *D. melanogaster* and the *simulans* clade are in different locations.** FISH image of mitotic chromosomes showing *Rsp-like* (red) and *1.688* (green) satellites. Chromosomes are counterstained using DAPI.

**Figure S2. Fluorescence in-situ hybridization of *Rsp-like* and *1.688* satDNA probes to *D. simulans* polytene chromosome.** *Rsp-like* is labeled in green, *1.688* in red. Chromosomes are counterstained using DAPI. This figure shows that *1.688* and *Rsp-like* are distributed throughout the X chromosome euchromatin with a higher prevalence of these repeats in the middle of the X chromosome visible in polytenes. This is consistent with the distribution in our assemblies.

**Figure S3. Overview of satDNA dynamics on X chromosome.** chrX: represents the X chromosome assembly of *D. melanogaster* with numbers 0–23 indicating position in Mb. *Rsp-like* clusters with more than 3 monomers are shown as black, red, blue, or green vertical lines according to species. 'Dead' clusters (cluster with ≤ 3 monomers) are shown by more transparent vertical lines. The position of *1.688* clusters for each species is shown by small gray dots below the *Rsp-like* lines for each species.

**Figure S4. Distribution of euchromatic satDNA cluster sizes on X chromosome.** Each satellite cluster in X chromosome euchromatin is represented by a black dot. 'Size of locus' along the y-axis indicates the number of repeat copies in a given cluster. For cases in which multiple clusters of the same size are present, black dots spread horizontally and appear as a horizontal black line. Box color indicates repeat family (blue for *Rsp-like,* orange for *1.688*).

**Figure S5. Nearest sequence elements to *1.688* satDNA clusters.** Each dot represents the nearest element to a single satellite cluster. The x-axis indicates the nearest neighboring element upstream and downstream; points are spread out along the x-axis in order to resolve them. The y-axis indicates the number of bases between a satellite cluster and its nearest neighbor with positive and negative values indicating nearest upstream elements and downstream elements respectively.

**Figure S6. Nearest sequence elements to *Rsp-like* satDNA clusters.** See caption of Fig. S5 for explanation.

**Figure S7. Phylogenetic relationships among *1.688* repeats in *Drosophila mauritiana*.** Each terminal represents an individual repeat monomer from the X chromosome. Colored tip terminals indicate euchromatic repeats with tips of the same color indicating repeats from the same cytoband. Grey tip terminals represent repeats from heterochromatic loci (defined as unassigned scaffolds in the assembly), which we subsetted to include only unique variants. The black tip terminal indicates the outgroup, which is a consensus sequence of X chromosome euchromatic loci for either *1.688* or *Rsp-like* from *D. erecta*, a sister species to *D. melanogaster* + the *simulans* clade. Tip size is scaled to reflect proportion of eccDNA reads mapping to a given variant, expressed as reads-per-million (RPM) / frequency of variant. Black rectangles indicate nodes with a bootstrap support ≥ 90. Maximum likelihood trees were inferred in RAxML with nodal support calculated following 100 bootstrap replicates. Branch length is shown proportional to relative divergence.

**Figure S8. Phylogenetic relationships among *Rsp-like* repeats in *Drosophila mauritiana*.** See caption S6 for explanation.

**Figure S9. Phylogenetic relationships among *1.688* repeats in *Drosophila sechellia*.** See caption S6 for explanation.

**Figure S10. Phylogenetic relationships among *Rsp-like* repeats in *Drosophila sechellia*.** See caption S6 for explanation.

**Figure S11. Phylogenetic relationships among *1.688* repeats in *Drosophila simulans*.** See caption S6 for explanation.

**Figure S12. Phylogenetic relationships among *Rsp-like* repeats in *Drosophila simulans*.** See caption S6 for explanation.

**Figure S13. Phylogenetic relationships among *1.688* repeats in *Drosophila melanogaster*.** See caption S6 for explanation.

**Figure S14. Phylogenetic relationships among *Rsp-like* repeats in *Drosophila melanogaster*.** See caption S6 for explanation.

**Figure S15. Phylogenetic relationship of *1.688* repeats across all study species part A.** Euchromatic repeats from all species are included in the analysis with branch color indicating species identity. See the caption for Fig. S7 for additional explanation.

**Figure S16. Phylogenetic relationship of *1.688* repeats across all study species part B.** See caption S14 for explanation.

**Figure S17. Phylogenetic relationship of *Rsp-like* repeats across all study species part A.** See caption S14 for explanation.

**Figure S18. Phylogenetic relationship of *Rsp-like* repeats across all study species part B.** See caption S14 for explanation.

**Figure S19. Age of satDNA clusters as inferred by pattern of gene conversion.** Distribution of distance metric dY for clusters of >3 repeats. The red dotted line indicates the cutoff for new vs older cluster (0.05). See text for description of dY metric.

**Figure S20. dY scatter plots.** Each circle represents a repeat cluster of >3 monomers, with the circle size scaled to reflect the cluster size. First-center and last-center indicates the average genetic distance between the first/last repeat in the cluster and the middle repeats. Middle-middle indicates average genetic distance between repeats in the middle of the cluster. We define a metric for age of a cluster, dY, as the maximum of first-middle vs middle-middle and last-middle vs middle- middle.

**Figure S21. Hi-C** Circle plot similar to showing interactions of sequences 10 Kb windows of flanking *1.688* repeats across the X chromosome. Lines connecting points on the circumference of the circle show connections between 10 Kb windows with normalized interaction counts greater than 40. We plotted interactions of flanking windows rather than the *1.688* sequences themselves to avoid issues related to low mappability of repetitive sequences.

**Figure S22. Sequence similarity across the X chromosome of *Rsp-like* and adjacent *1.688* repeats.** Plot shows repeats of high sequence similarity within and across cytobands 1–12 for *Rsp-like* repeats, and any *1.688* repeats that are in clusters adjacent (within 100 bases) to *Rsp-like* clusters. Each point around the circumference of the circle represents a single repeat. The position of *1.688* repeats are shown by orange dots, all remaining positions represent *Rsp-like* repeats; both repeat types are shown in the relative order in which they occur in cytobands 1–12 shown by colored bars. Lines are drawn between any repeats showing greater than 95% sequence similarity.

**Figure S23. Verification of linear DNA digest and presence of eccDNA in *D. melanogaster.*** We used 2D gel analysis to confirm/show the presence of *1.688* and *Rsp* eccDNA in *D. melanogaster.* A) PCR analysis of *D. melanogaster* (iso 1) genomic DNA before and after digestion with exoV. The rp49 and tRNA results (top) indicate a significant loss of linear DNA while the mitoCOI results (bottom) indicate little loss of circular DNA during exoV digestion. Consistent with a previous report (Cohen, et al. 2003), the results indicate that sequences derived from 5S, histone 2A, and *1.688* tandem repeats are present on extrachromosomal circles. PCR product after exoV digestion also suggests that extrachromosomal circles are derived from *Rsp* repeats. (B) 2D gel analysis to confirm *Rsp* extrachromosomal circles. The diagram at the top details gel electrophoresis and where double stranded linear DNA, open DNA circles, and single stranded DNA migrate. The middle gel was southern blotted and probed with *Rsp*. The bottom gel was southern blotted and probed with *1.688*. Both blots show hybridization to slow migrating DNA which has been attributed to open circles (Cohen and Lavi 1996; Cohen, et al. 1999). Genomic DNA used in the blots was isolated from Raleigh 370- a strain with ~9 fold more *Rsp* repeats than iso 1.

**Figure S24. Verification of linear DNA digestion in *simulans* clade and *D. melanogaster.*** PCR analysis of the genomic DNAs used for sequencing before and after digestion with exoV. The mitochondria cytochrome C oxidase results indicate the circular mitochondrial genome is present at a comparable level in both digested and undigested samples (dilution series in black at top). The rp49 results indicate a significant loss of linear DNA after exoV digestion (dilution series in gray at bottom).

**Fig. S25. Abundance of eccDNA for repeat cateogories in all four species.** Repeats in the genome are categorized into Satellite, LTR retrotransposon, non-LTR retrotransposon, DNA transposon and rolling-circle (RC) transposon. Boxplots summarize log-scale RPM values and the log scale values of repeats in each category.

**Figure S26. Histograms of genomic attributes showing highly significant association with top 25 MEME motifs.** MEME identified 25 motifs that were abundant in sequences flanking *Rsp-like* clusters. Top 25 motifs were mapped to the X chromosome, and genomic attributes containing each motif were ranked based on the significance of their association with that motif following Bonferroni correction of p-values. Genomic attributes are listed along the x-axis. The y-axis reports the number of top 25 MEME motifs showing highly significant association with a given genomic attribute. A genomic attribute with a y-value of 10 indicates that for 10 of the top 25 meme motifs, that genomic attribute ranked in the top 1% of most significant p-values.

#### SUPPLEMENTARY FIGURES

Fig. S1

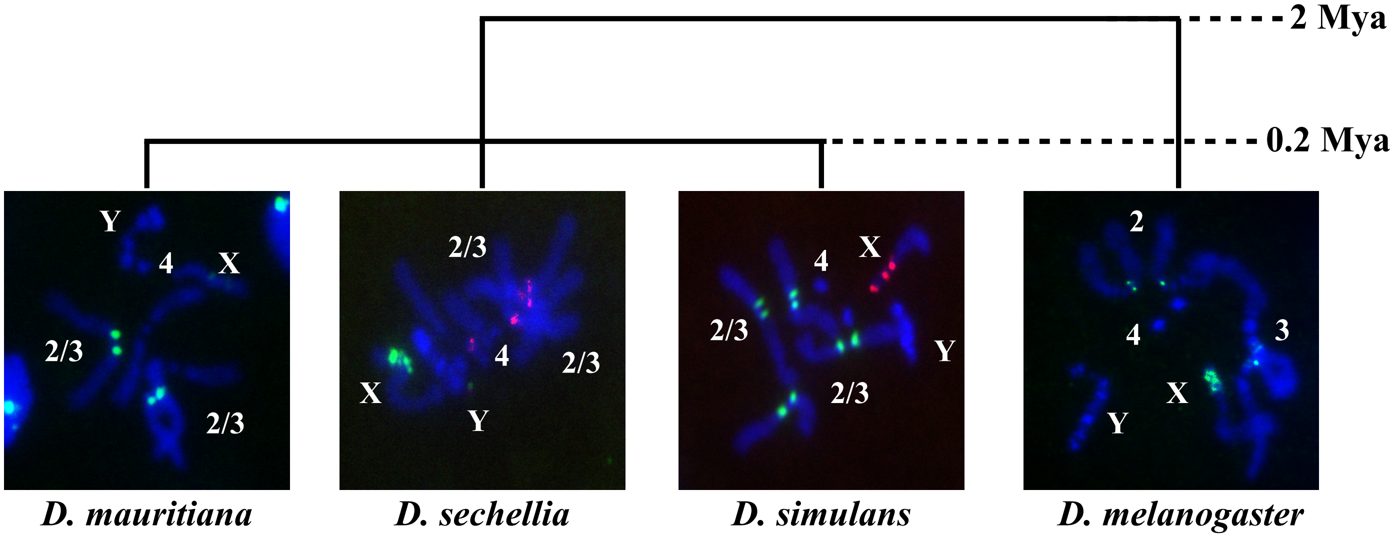
2 Mya
0.2 Mya
Y
4
X
2/3
2/3
2/3
X
4
Y
2/3
4
X
2/3
Y
2/3
2
4
3
X
Y
D. mauritiana
D. sechellia
D. simulans
D. melanogaster

Fig. S2

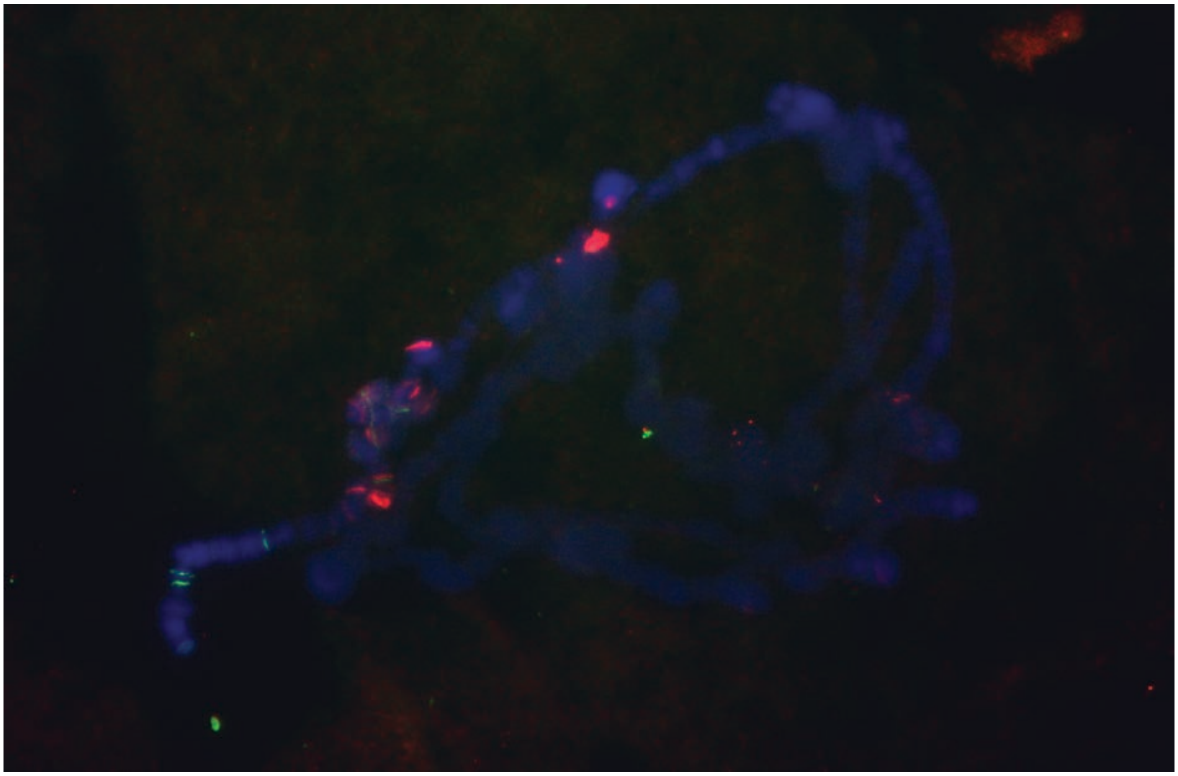

Fig. S3

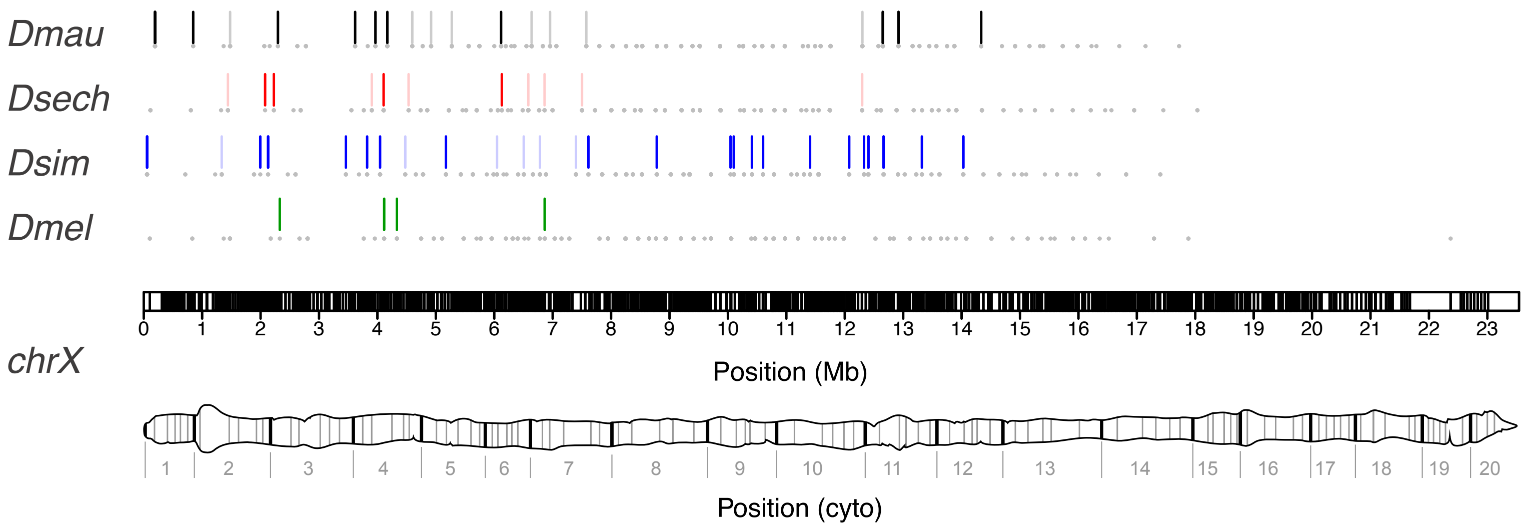
Dmau
Dsech
Dsim
Dmel
chrX
0 1 2 3 4 5 6 7 8 9 10 11 12 13 14 15 16 17 18 19 20 21 22 23
Position (Mb)
1 2 3 4 5 6 7 8 9 10 11 12 13 14 15 16 17 18 19 20
Position (cyto)

Fig. S4

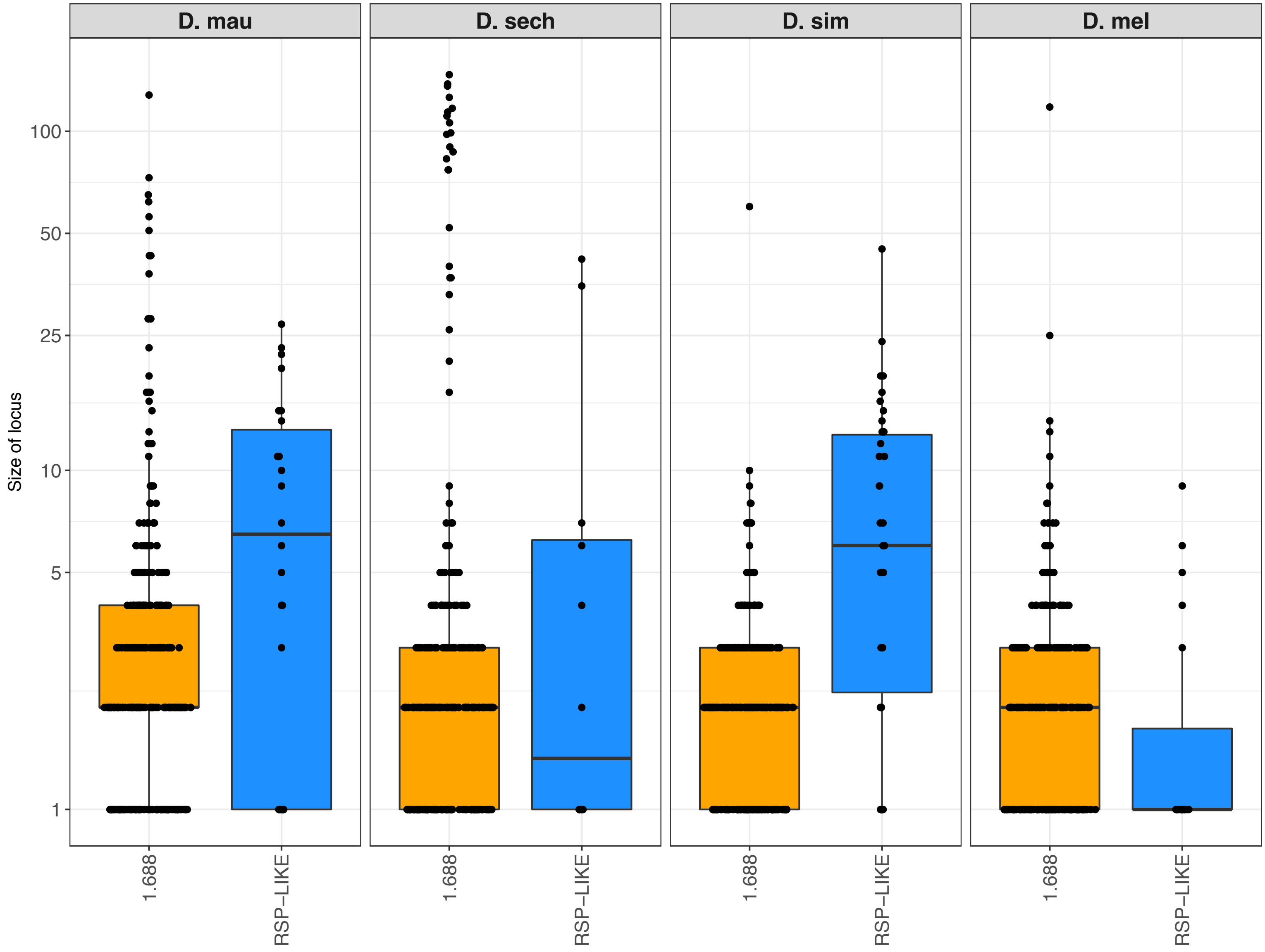

D. mau
D. sech
D. sim
D. mel
Size of locus
100
50
25
10
5
1
1.688
RSP–LIKE
1.688
RSP–LIKE
1.688
RSP–LIKE
1.688
RSP–LIKE

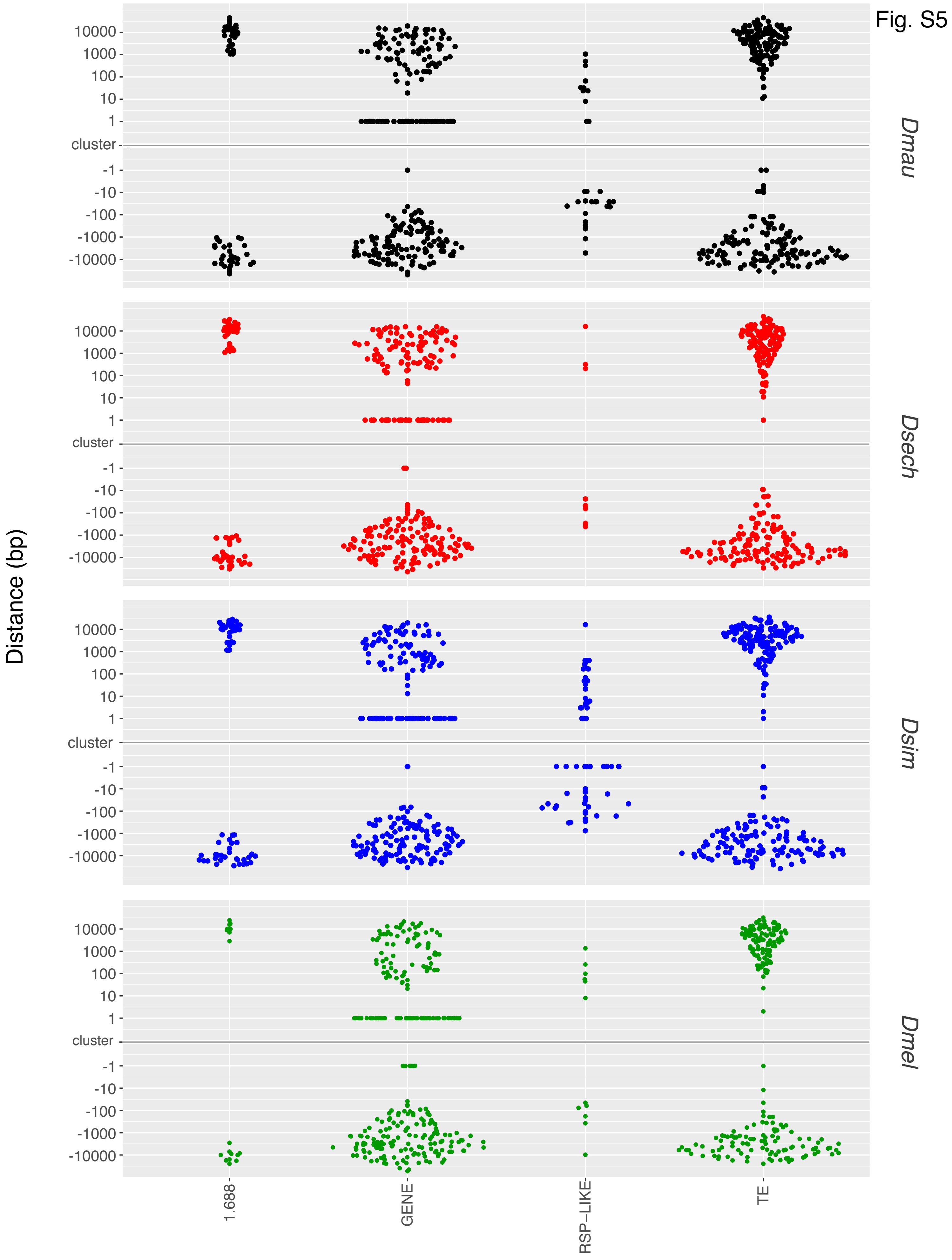
Fig. S5
Distance (bp)
10000
1000
100
10
1
cluster
-1
-10
-100
-1000
-10000
Dmau
Dsech
Dsim
Dmel
1.688
GENE
RSP–LIKE
TE

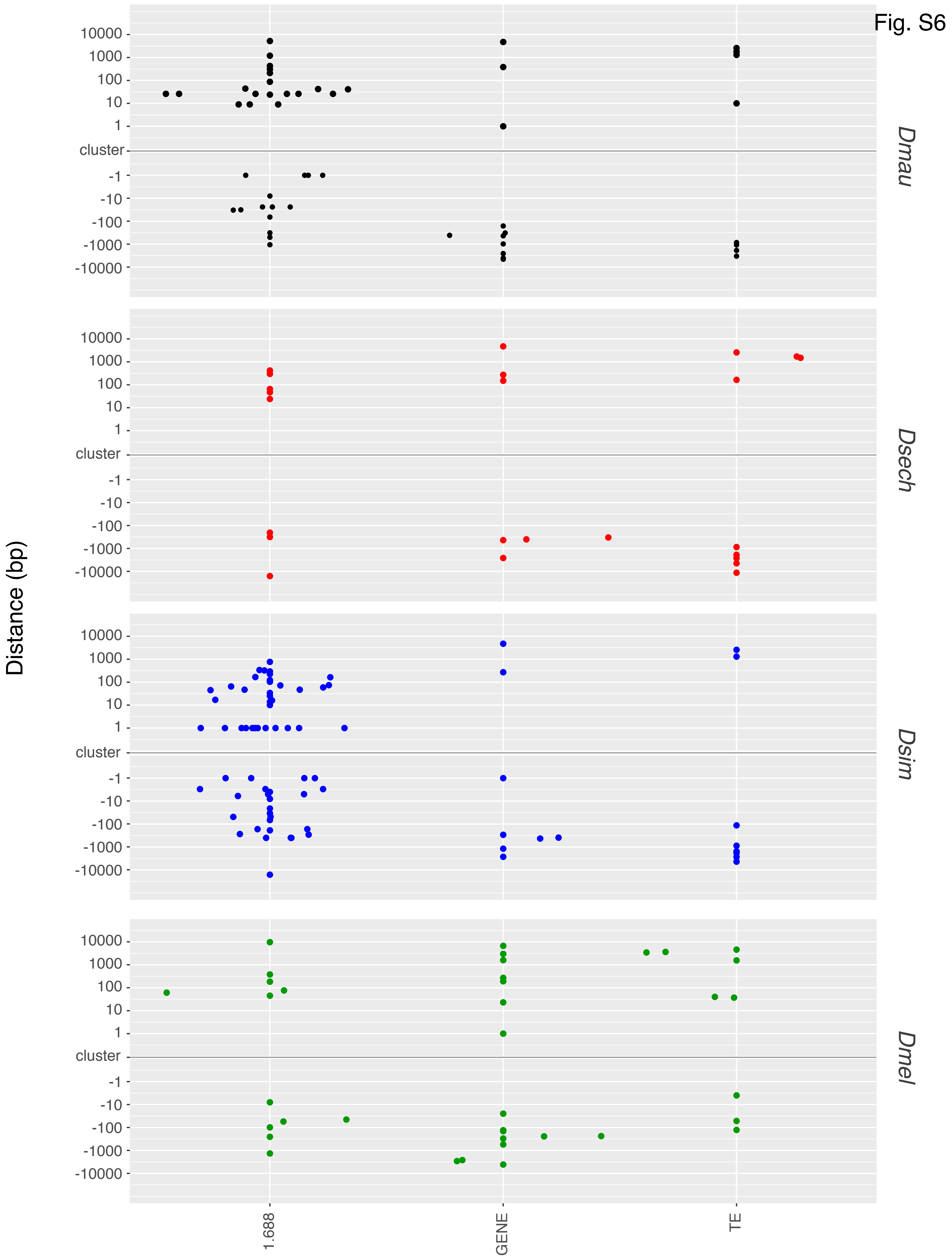
Fig. S6
Distance (bp)
10000
1000
100
10
1
cluster
-1
-10
-100
-1000
-10000
Dmau
Dsech
Dsim
Dmel
1.688
GENE
TE

Fig. S7

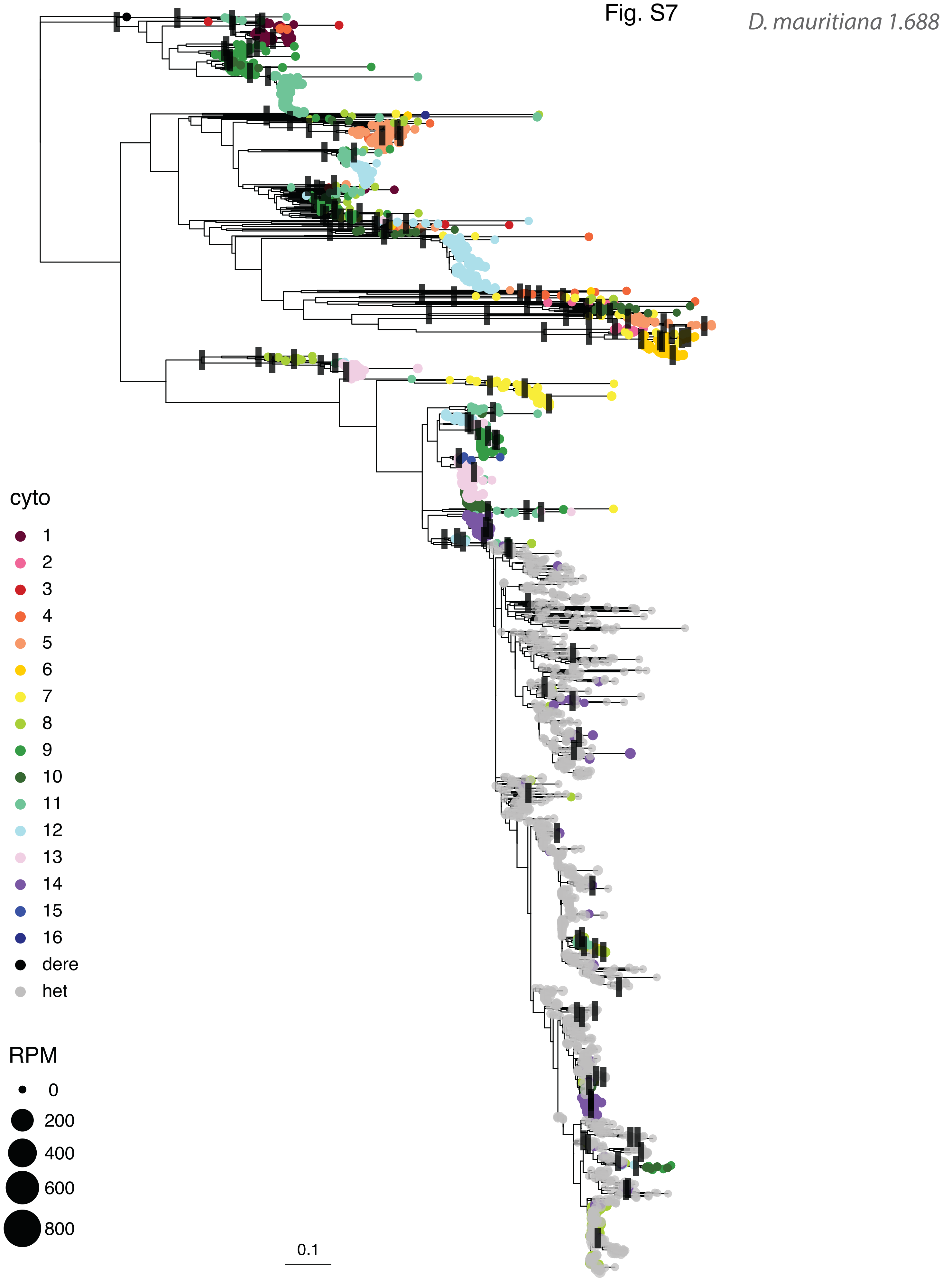
D. mauritiana 1.688
cyto
dere
het
RPM
0
200
400
600
800
0.1

Fig. S8

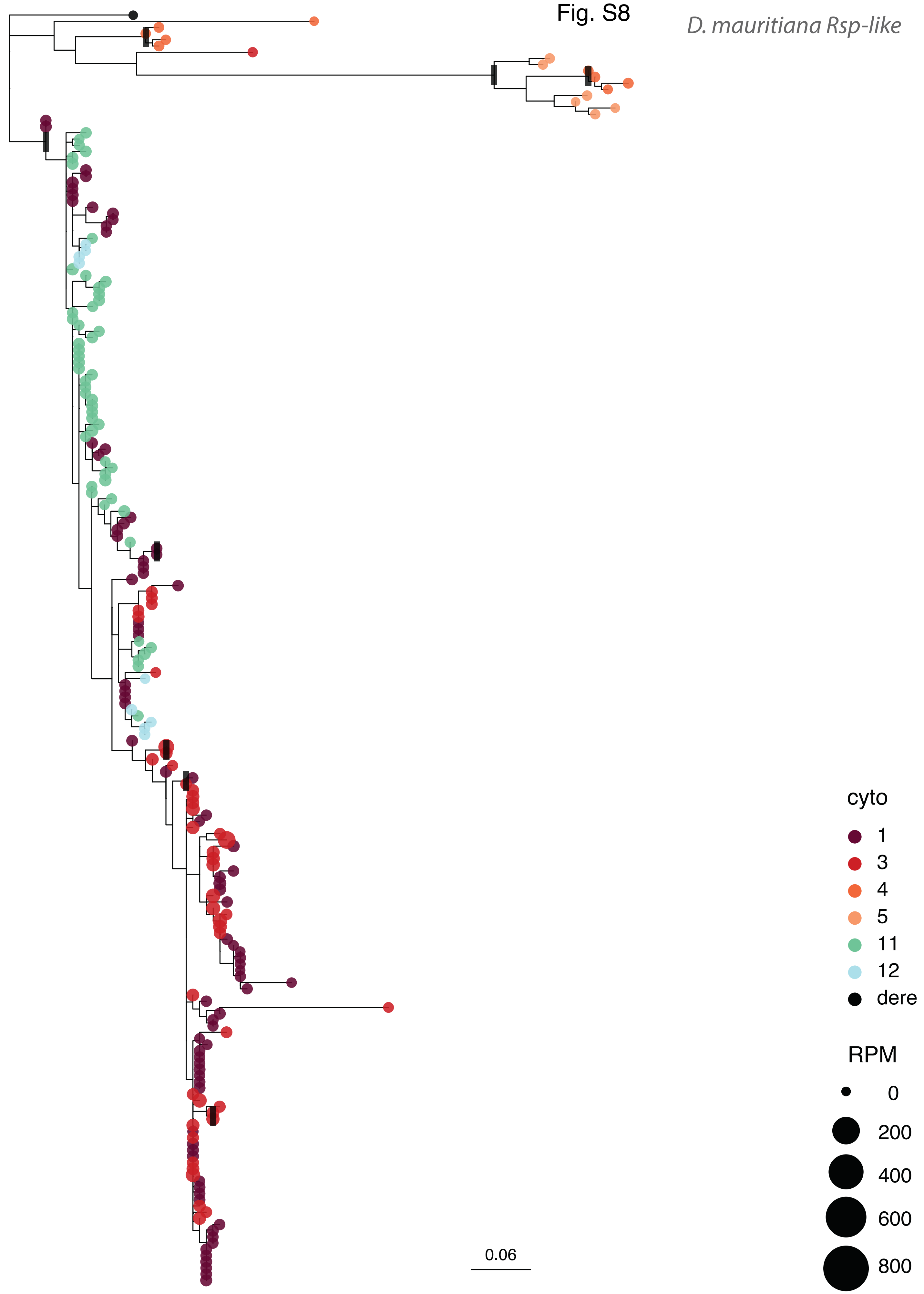
D. mauritiana Rsp-like
cyto
1
3
4
5
11
12
dere
RPM
0
200
400
600
800
0.06

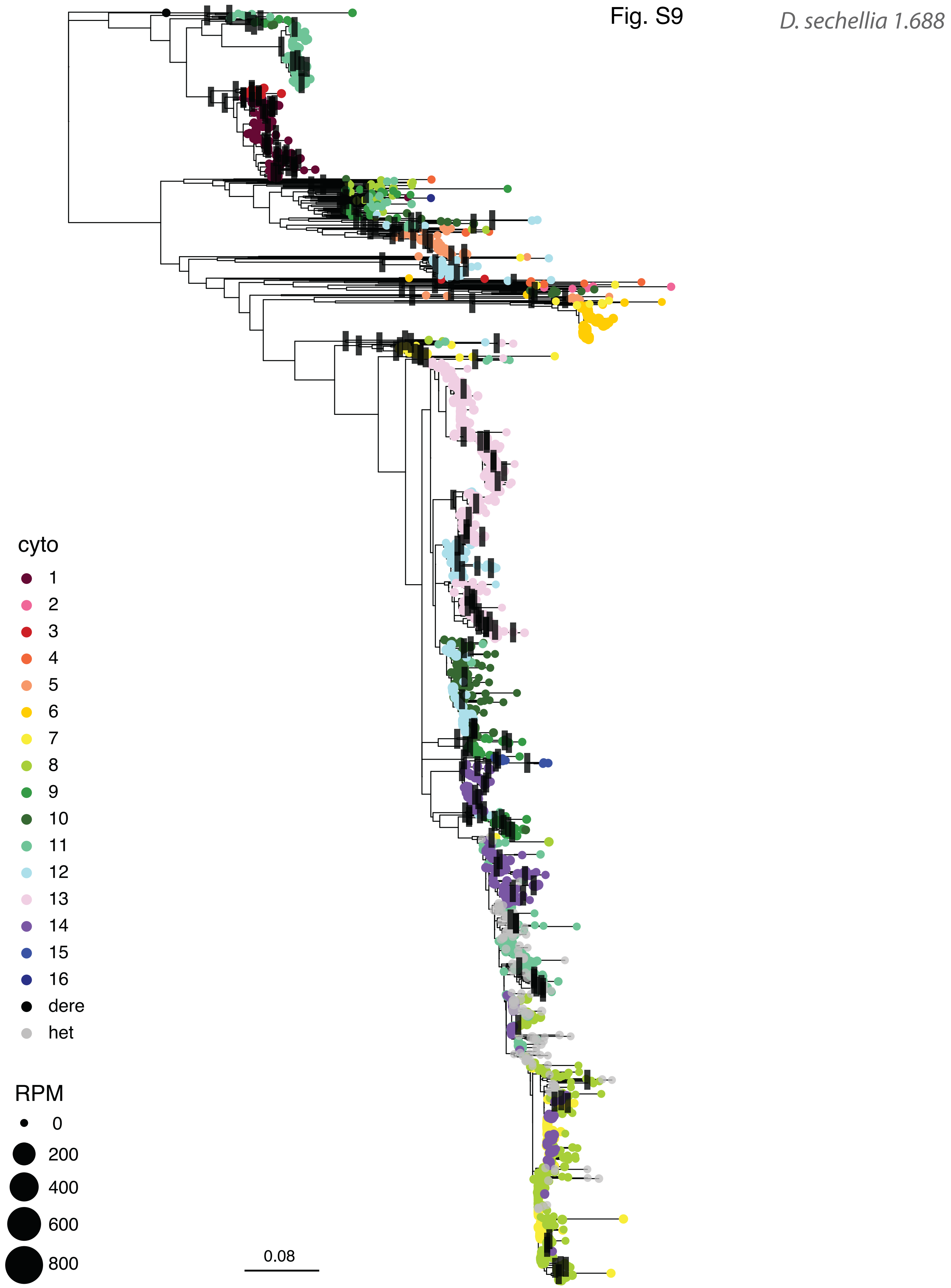
Fig. S9
D. sechellia 1.688
cyto
dere
het
RPM
0
200
400
600
800
0.08

Fig. S10

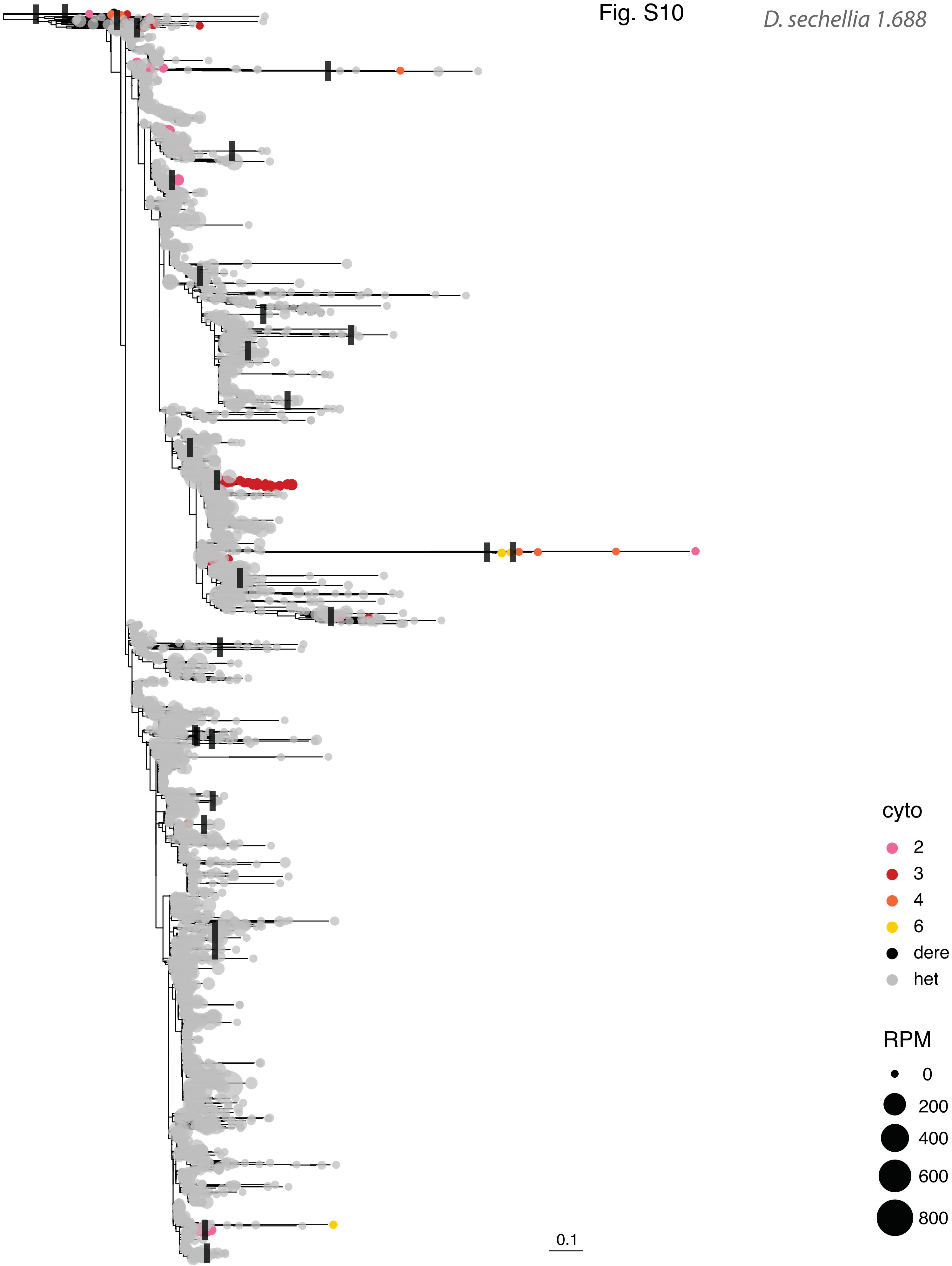

Fig. S11

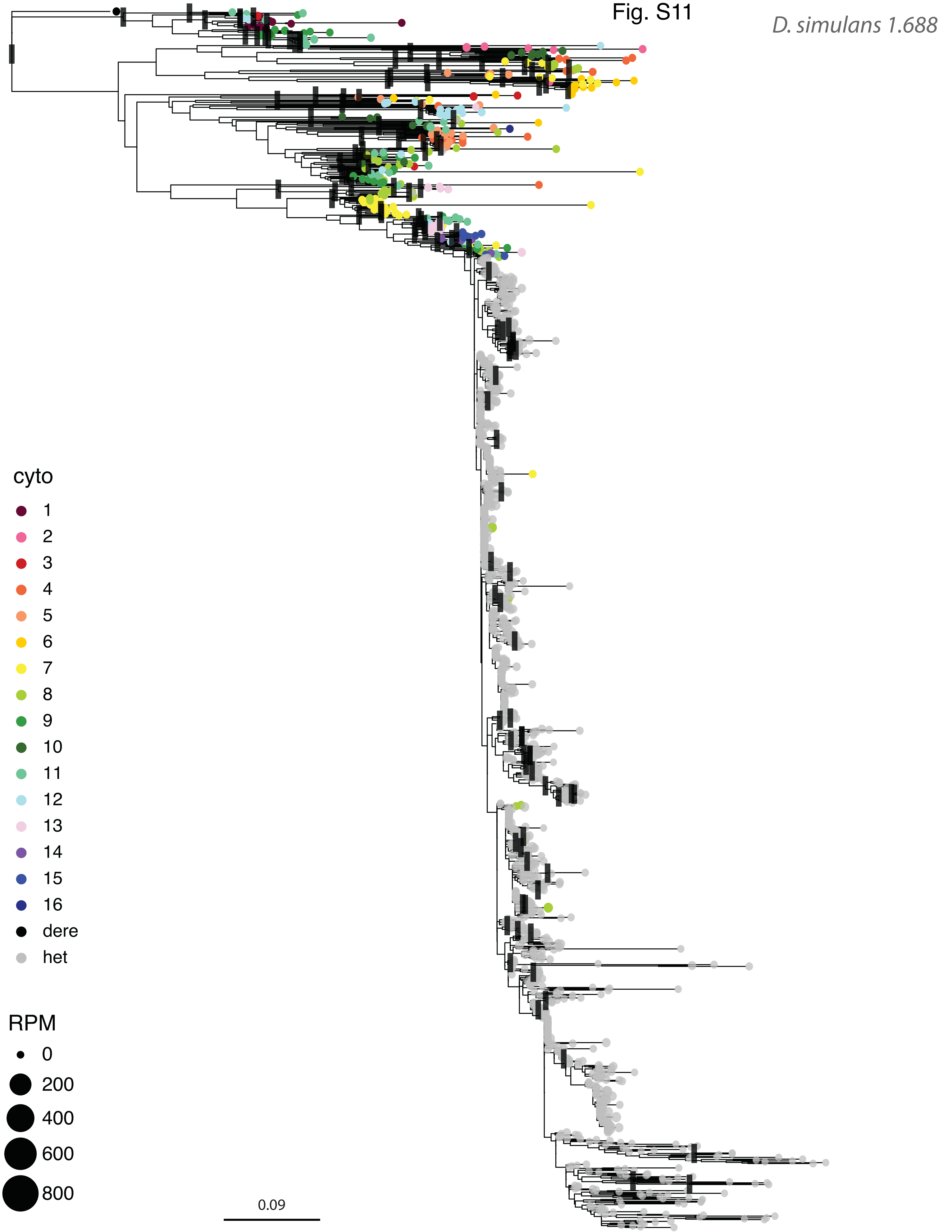
D. simulans 1.688
cyto
dere
het
RPM
0
200
400
600
800
0.09

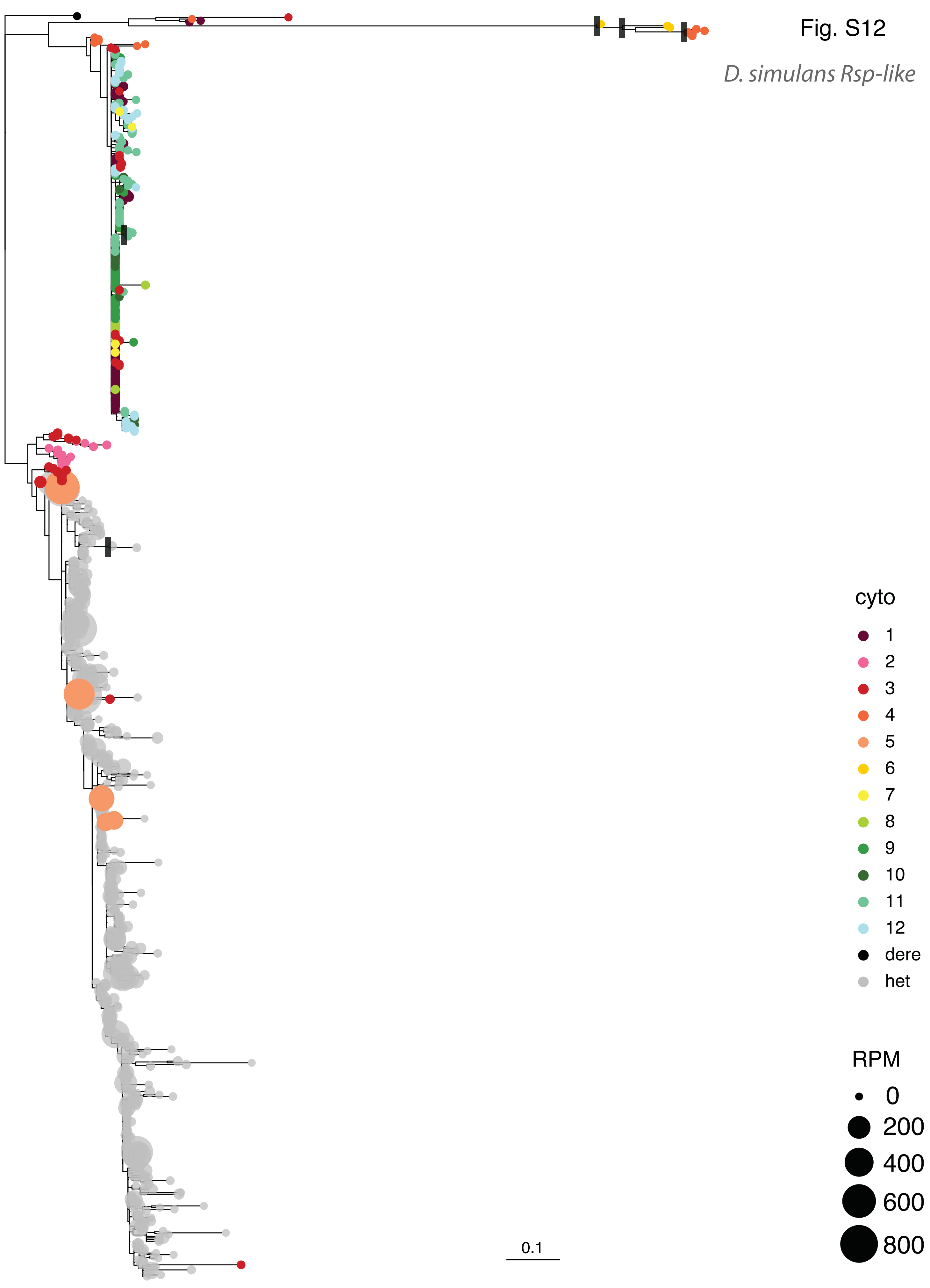
Fig. S12
D. simulans Rsp-like
cyto
1
2
3
4
5
6
7
8
9
10
11
12
dere
het
RPM
0
200
400
600
800
0.1

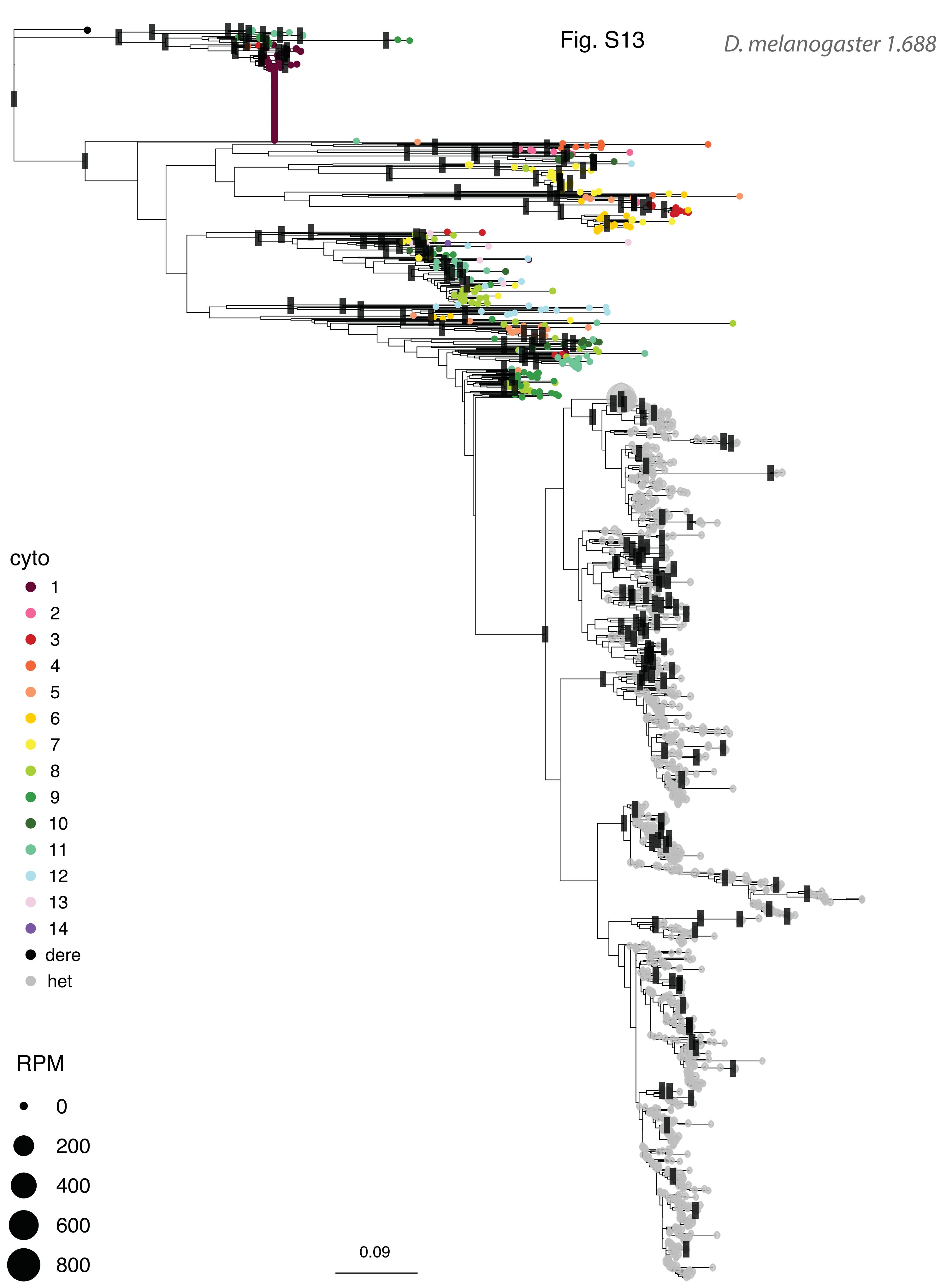
Fig. S13
D. melanogaster 1.688
cyto
1
2
3
4
5
6
7
8
9
10
11
12
13
14
dere
het
RPM
0
200
400
600
800
0.09

Fig. S14

*D. melanogaster Rsp-like*

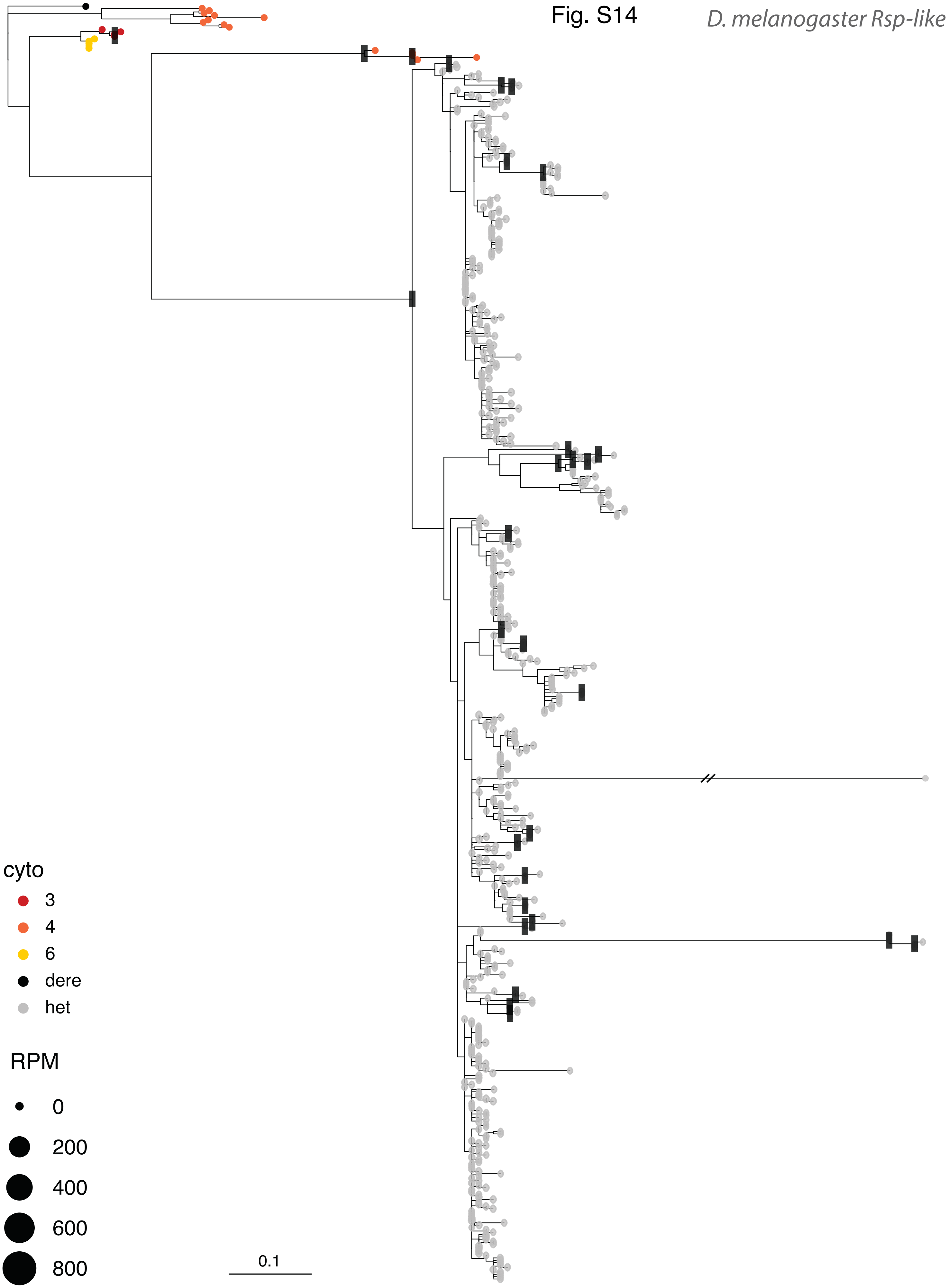

Fig. S15

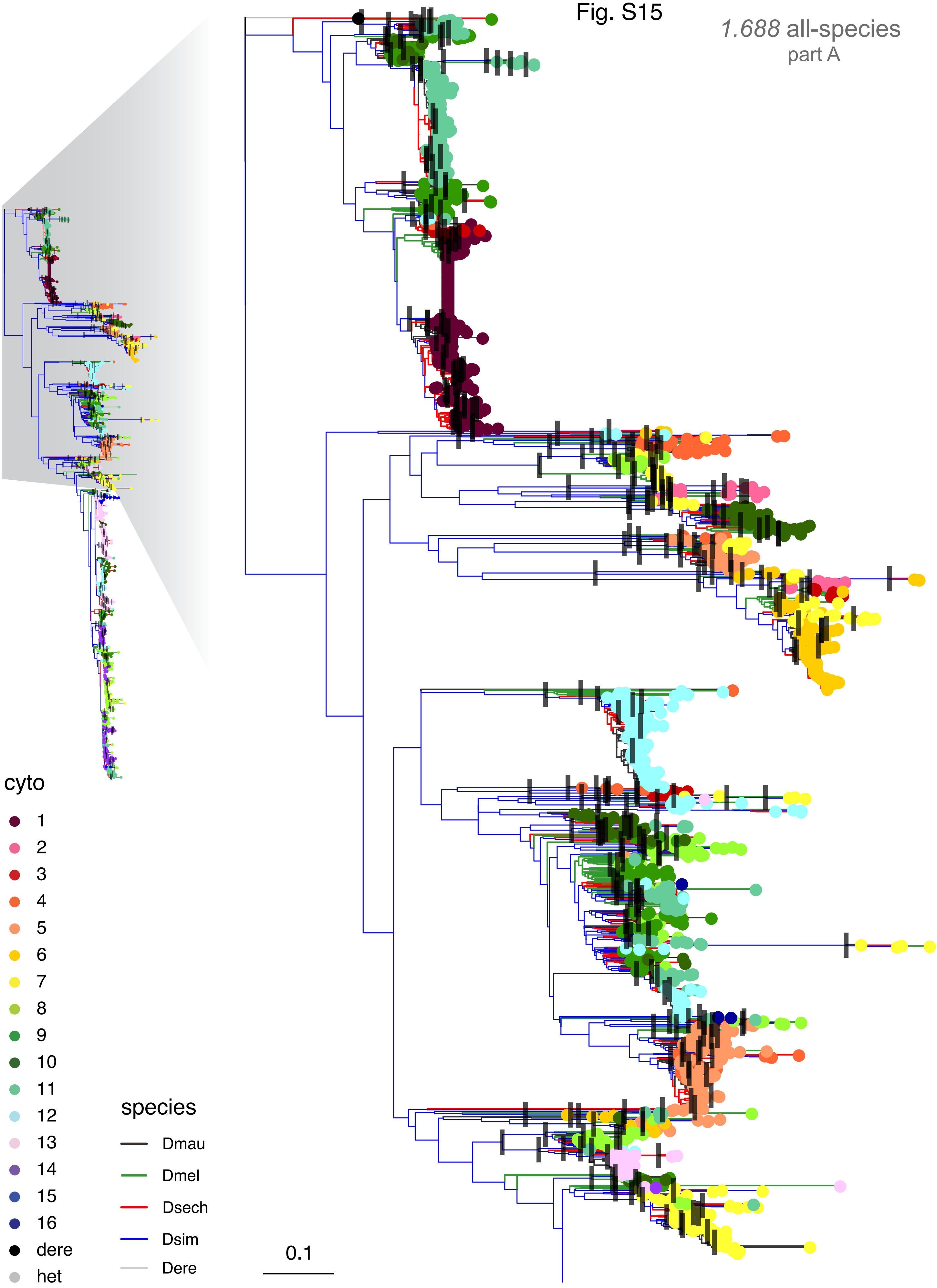

*1.688* all-species
part A
cyto
dere
het
species
Dmau
Dmel
Dsech
Dsim
Dere
0.1

Fig. S16

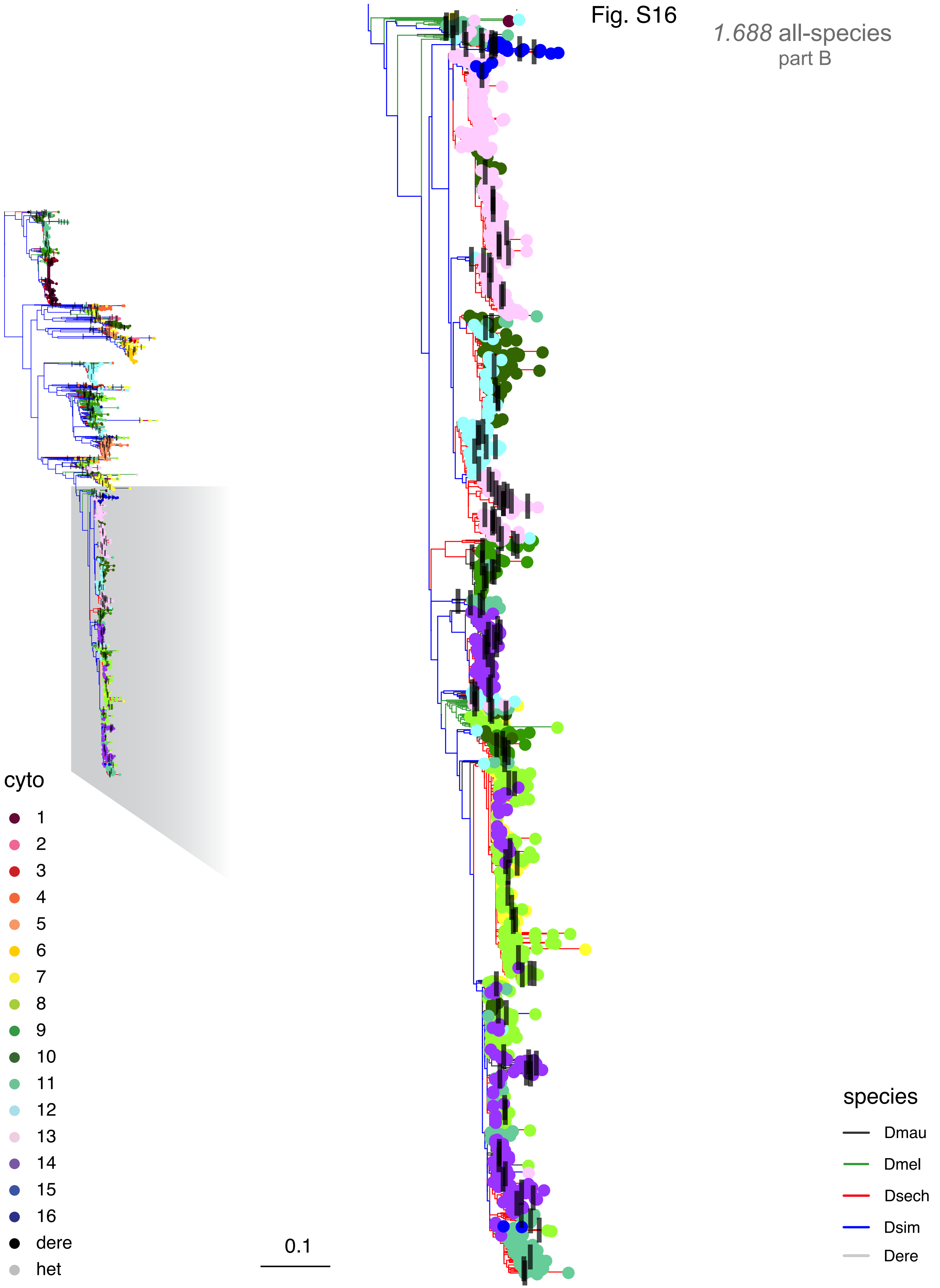
*1.688* all-species
part B
cyto
dere
het
0.1
species
Dmau
Dmel
Dsech
Dsim
Dere

Fig. S17

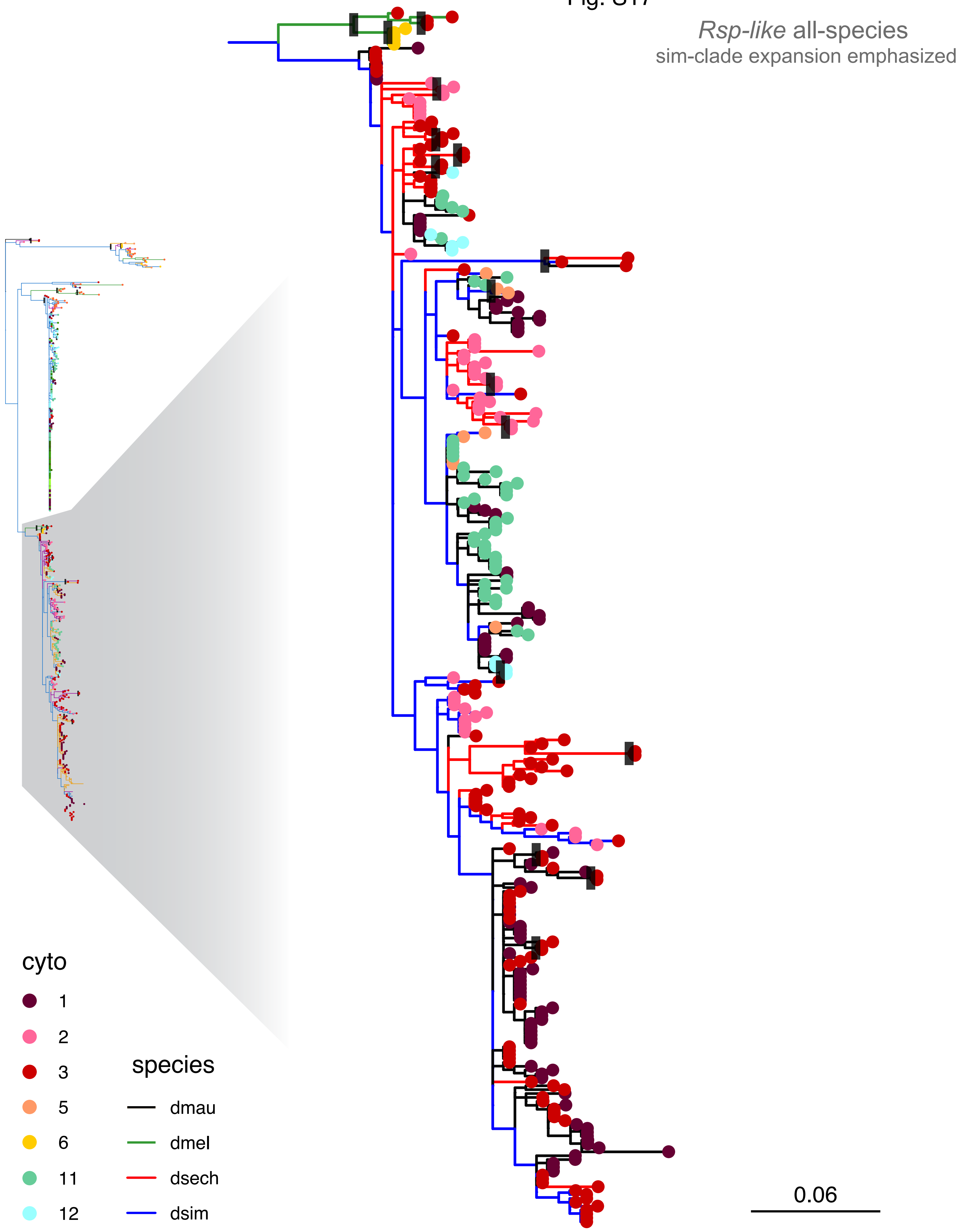

*Rsp-like* all-species
sim-clade expansion emphasized
cyto
1
2
3
5
6
11
12
species
dmau
dmel
dsech
dsim
0.06

Fig. S18

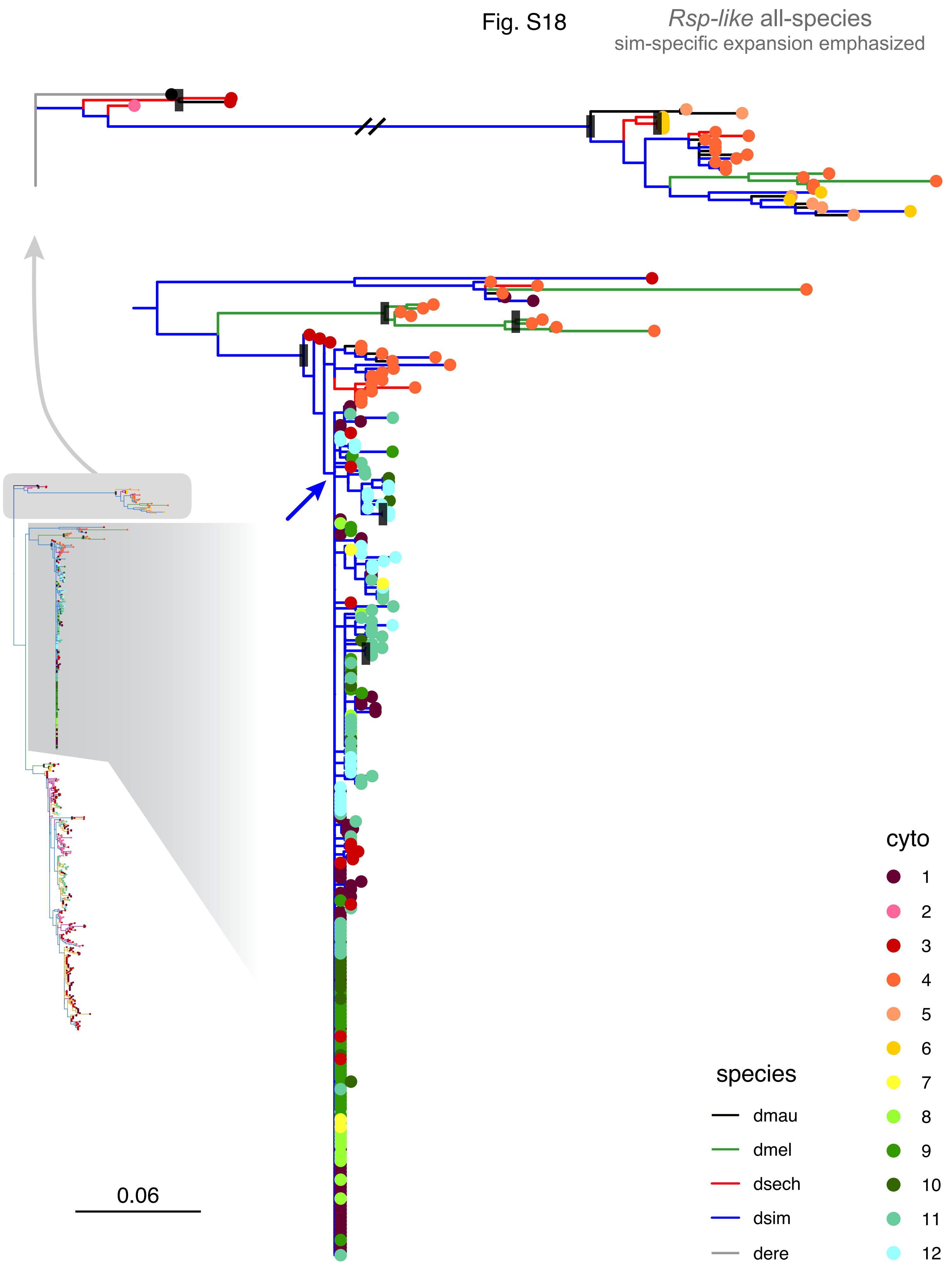
*Rsp-like* all-species
sim-specific expansion emphasized
0.06
cyto
1
2
3
4
5
6
7
8
9
10
11
12
species
dmau
dmel
dsech
dsim
dere

Fig. S19

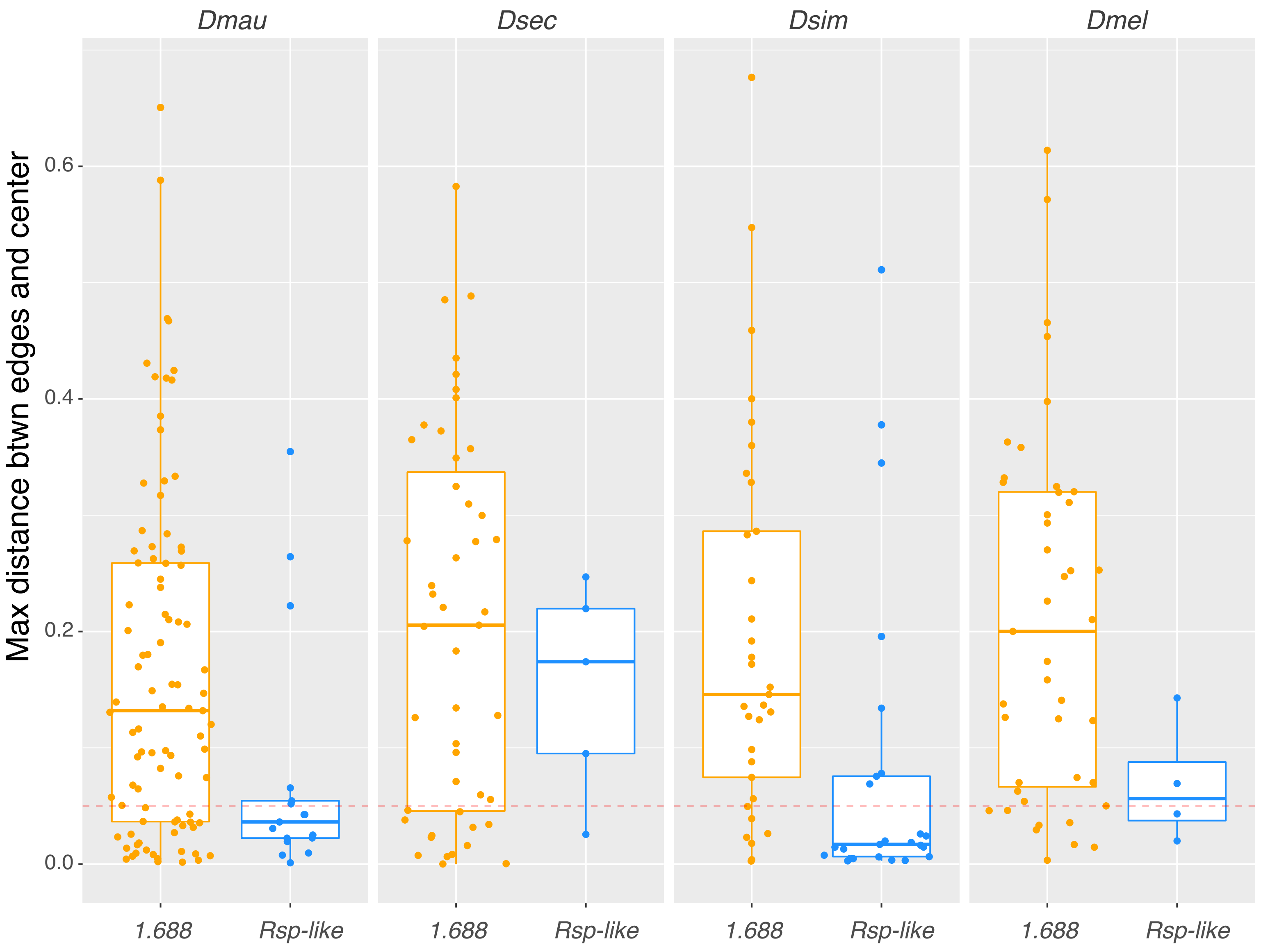

Dmau
Dsec
Dsim
Dmel
Max distance btwn edges and center
0.6
0.4
0.2
0.0
1.688
Rsp-like
1.688
Rsp-like
1.688
Rsp-like
1.688
Rsp-like

Fig. S20

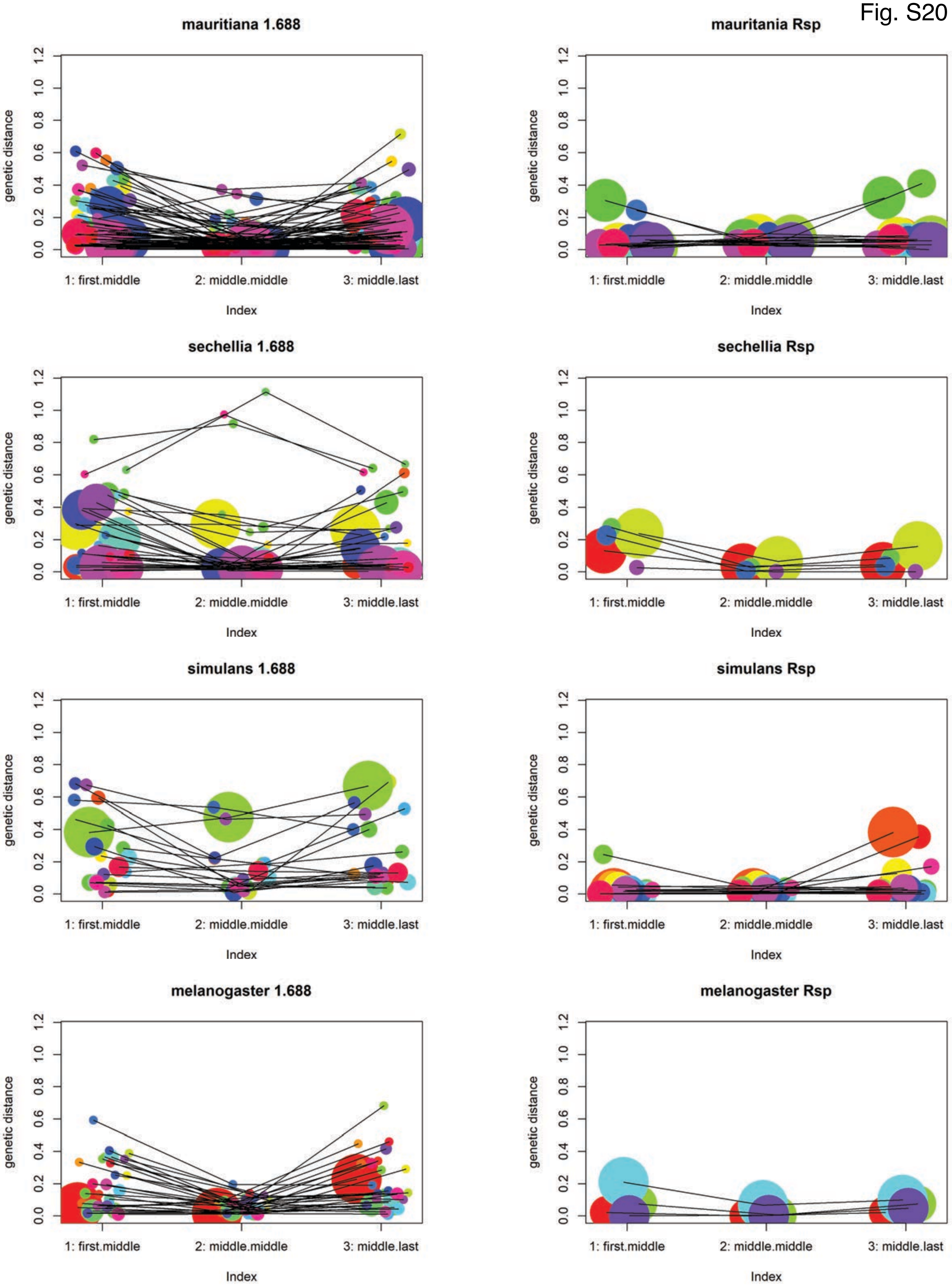

mauritiana 1.688
mauritania Rsp
sechellia 1.688
sechellia Rsp
simulans 1.688
simulans Rsp
melanogaster 1.688
melanogaster Rsp
genetic distance
0.0
0.2
0.4
0.6
0.8
1.0
1.2
1: first.middle
2: middle.middle
3: middle.last
Index

Fig. S21

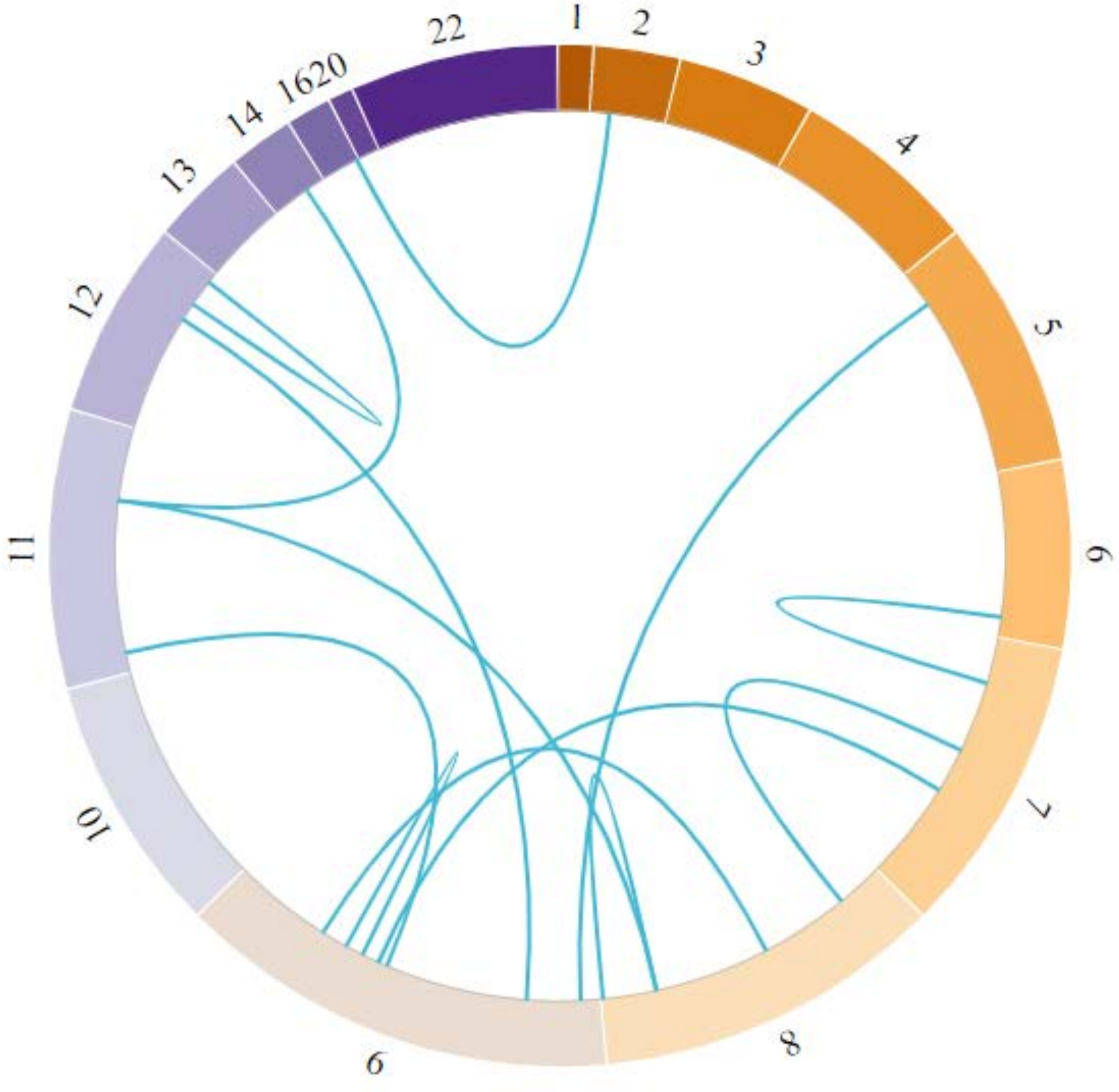

Fig. S22

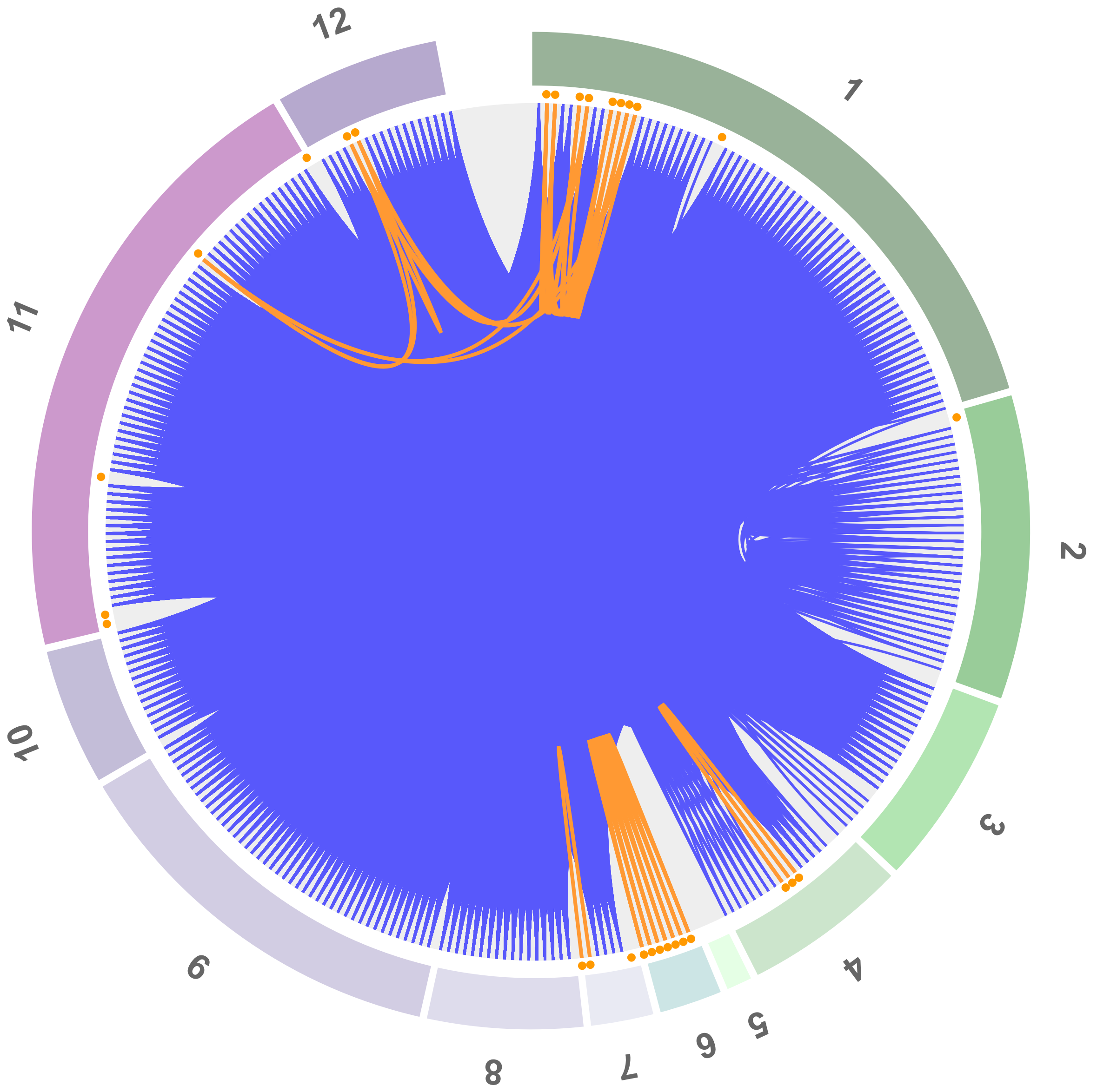

Fig. S23

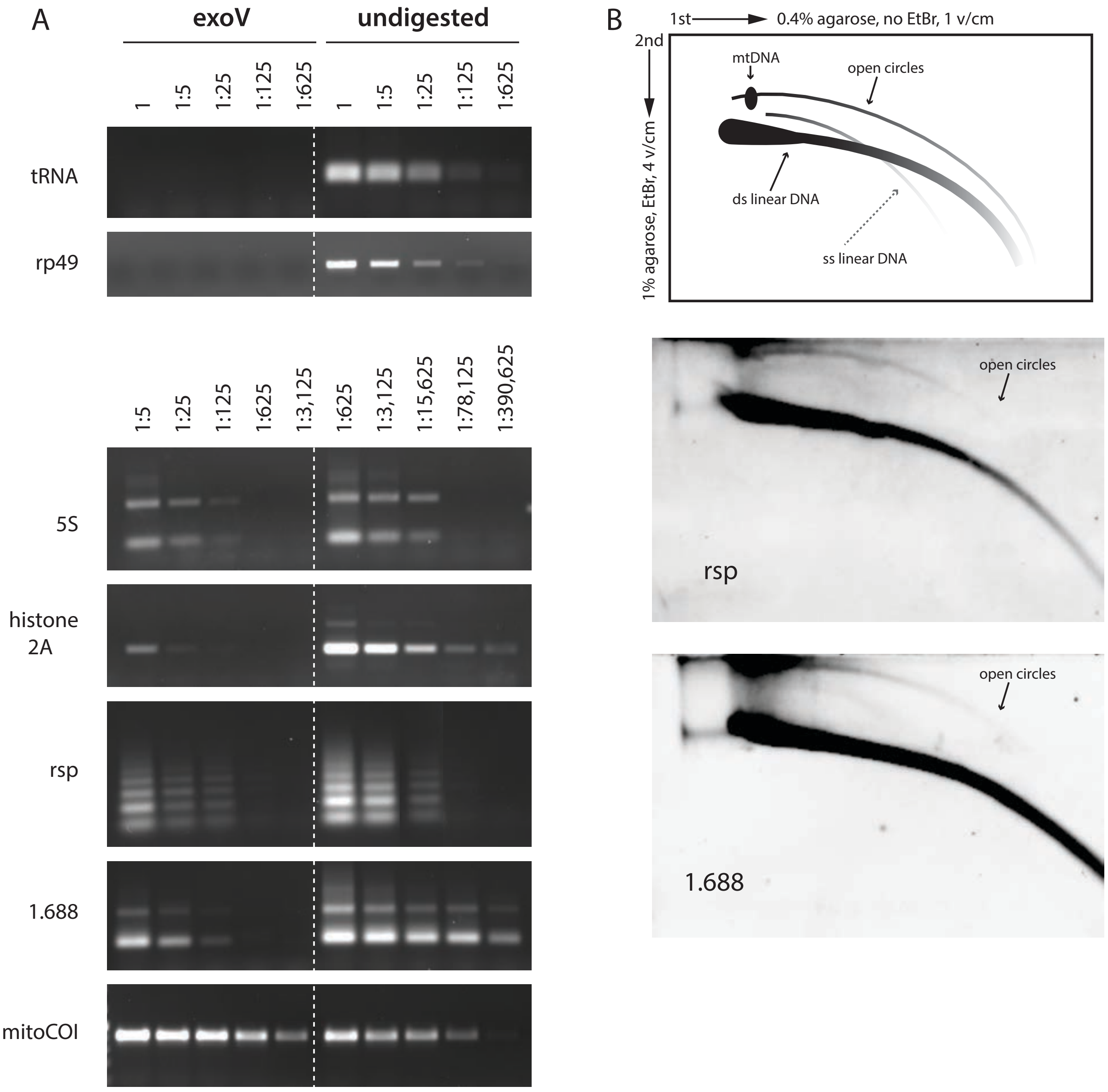
A
exoV
undigested
1
1:5
1:25
1:125
1:625
tRNA
rp49
1:625
1:3,125
1:15,625
1:78,125
1:390,625
5S
histone
2A
rsp
1.688
mitoCOI
B
1st
0.4% agarose, no EtBr, 1 v/cm
2nd
1% agarose, EtBr, 4 v/cm
mtDNA
open circles
ds linear DNA
ss linear DNA
open circles
rsp
open circles
1.688

Fig. S24

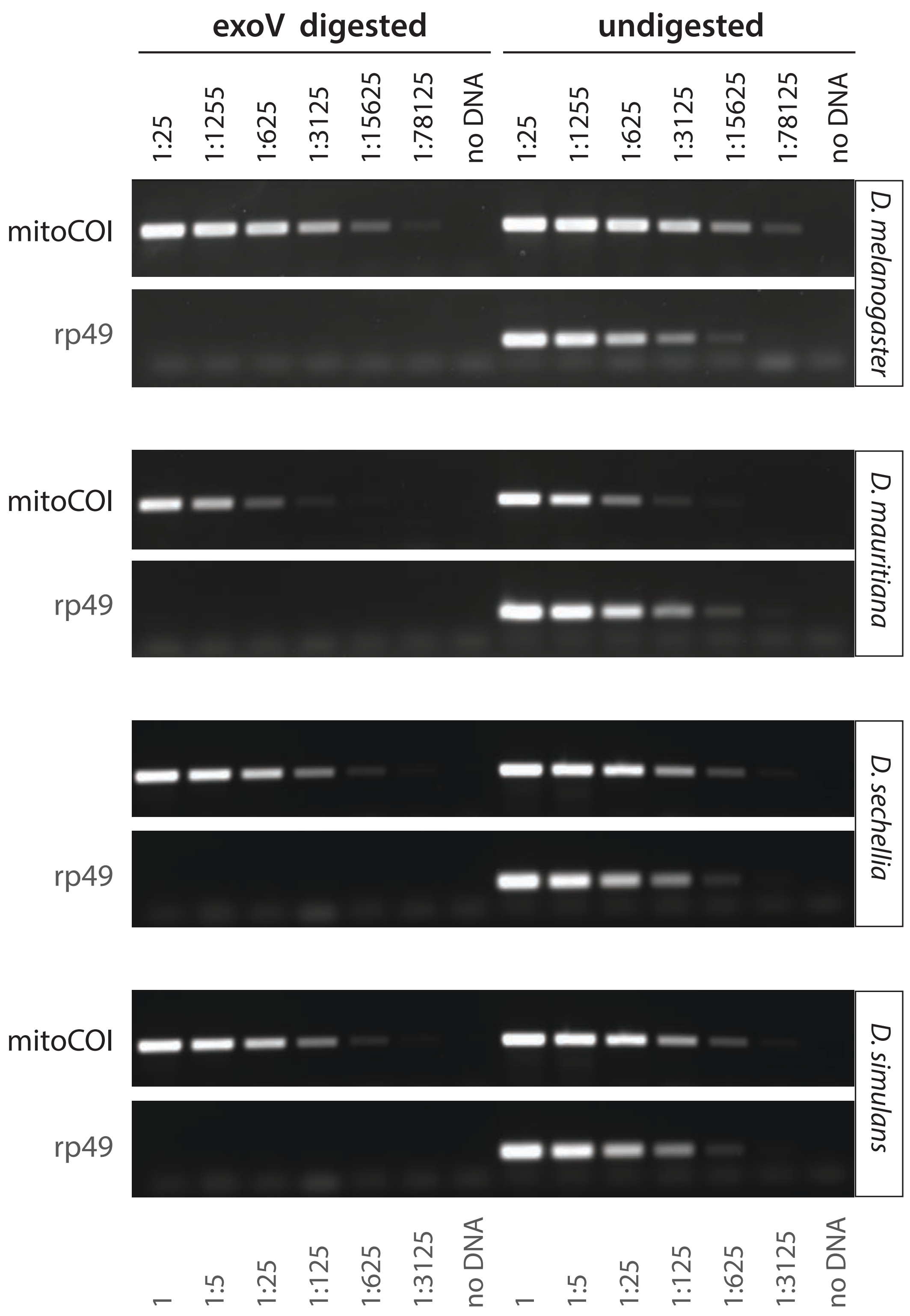

exoV digested
undigested
1:25
1:1255
1:625
1:3125
1:15625
1:78125
no DNA
mitoCOI
rp49
D. melanogaster
D. mauritiana
D. sechellia
D. simulans
1
1:5
1:125

Fig. S25

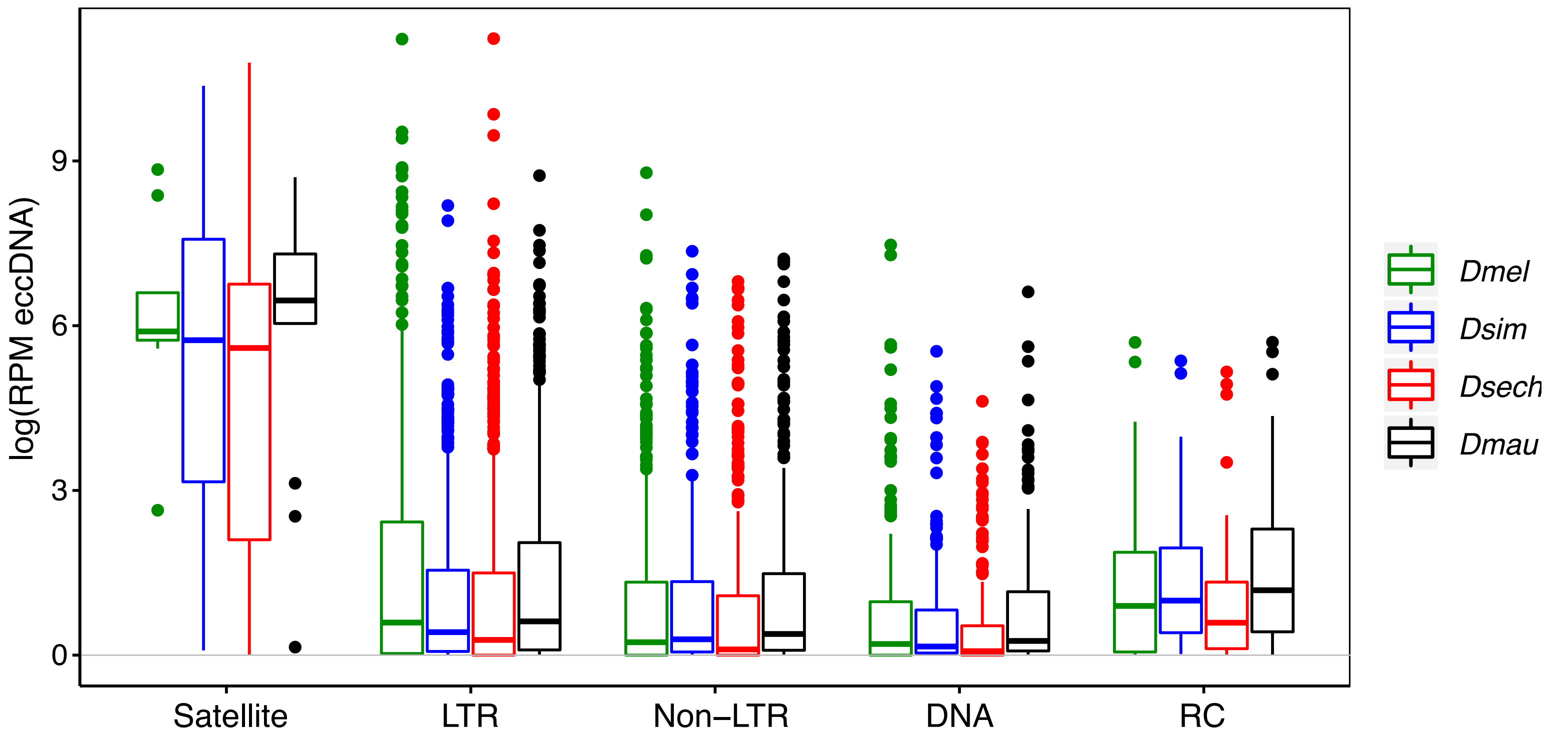

log(RPM eccDNA)
0
3
6
9
Satellite
LTR
Non−LTR
DNA
RC
Dmel
Dsim
Dsech
Dmau

Fig. S26

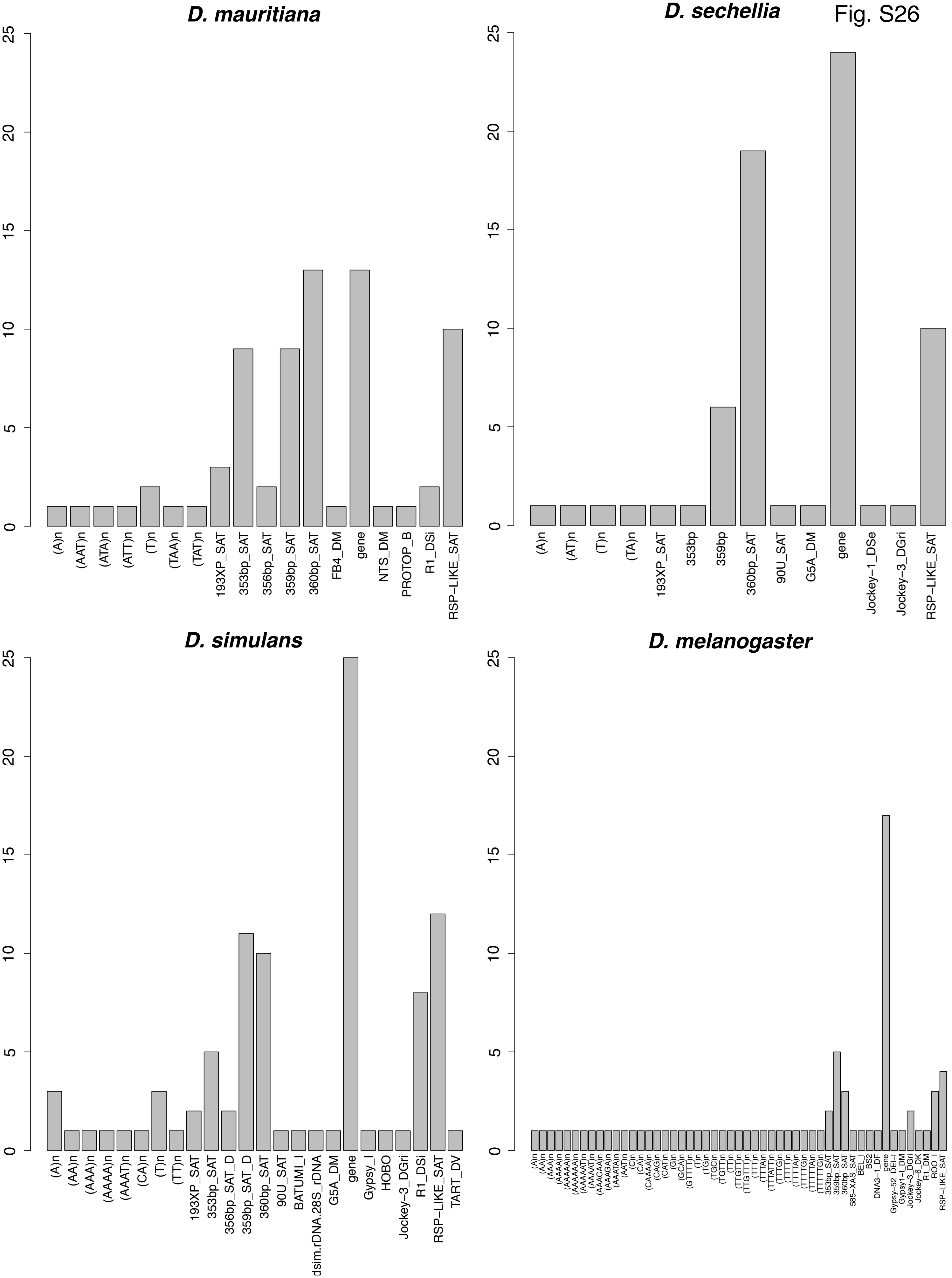

D. mauritiana
0
5
10
15
20
25
(A)n
(AAT)n
(ATA)n
(ATT)n
(T)n
(TAA)n
(TAT)n
193XP_SAT
353bp_SAT
356bp_SAT
359bp_SAT
360bp_SAT
FB4_DM
gene
NTS_DM
PROTOP_B
R1_DSi
RSP–LIKE_SAT
D. sechellia
(A)n
(AT)n
(T)n
(TA)n
193XP_SAT
353bp
359bp
360bp_SAT
90U_SAT
G5A_DM
gene
Jockey–1_DSe
Jockey–3_DGri
RSP–LIKE_SAT
D. simulans
(A)n
(AA)n
(AAA)n
(AAAA)n
(AAAT)n
(CA)n
(T)n
(TT)n
193XP_SAT
353bp_SAT
356bp_SAT_D
359bp_SAT_D
360bp_SAT
90U_SAT
BATUMI_I
dsim.rDNA.28S_rDNA
G5A_DM
gene
Gypsy_I
HOBO
Jockey–3_DGri
R1_DSi
RSP–LIKE_SAT
TART_DV
D. melanogaster
(A)n
(AA)n
(AAA)n
(AAAA)n
(AAAAA)n
(AAAAAA)n
(AAAAAT)n
(AAAAT)n
(AAACAA)n
(AAAGA)n
(AAATA)n
(AAT)n
(C)n
(CA)n
(CAAAA)n
(CAG)n
(CAT)n
(G)n
(GCA)n
(GTTTT)n
(T)n
(TG)n
(TGC)n
(TGTT)n
(TT)n
(TTGTT)n
(TTGTTT)n
(TTT)n
(TTTA)n
(TTTATT)n
(TTTG)n
(TTTT)n
(TTTTA)n
(TTTTG)n
(TTTTTA)n
(TTTTTG)n
353bp_SAT
359bp_SAT
360bp_SAT
585–XAS_SAT
BEL_I
BS2
DNA3–1_DF
gene
Gypsy–52_DEI–I
Gypsy1–I_DM
Jockey–3_DGri
Jockey–6_DK
R1_DM
ROO_I
RSP–LIKE_SAT
